# Improvements on marker-free images alignment for electron tomography

**DOI:** 10.1101/2020.05.22.110445

**Authors:** C.O.S. Sorzano, F. de Isidro-Gómez, E. Fernández-Giménez, D. Herreros, S. Marco, J.M. Carazo, C. Messaoudi

## Abstract

Electron tomography is a technique to obtain three-dimensional structural information of samples. However, the technique is limited by shifts occurring during acquisition that need to be corrected before the reconstruction process. In 2009, we proposed an approach for post-acquisition alignment of tilt series images. This approach was marker-free, based on patch tracking and integrated in free software. Here, we present improvements to the method to make it more reliable, stable and accurate. In addition, we modified the image formation model underlying the alignment procedure to include different deformations occurring during acquisition. We propose a new way to correct these computed deformations to obtain reconstructions with reduced artifacts. The new approach has demonstrated to improve the quality of the final 3D reconstruction, giving access to better defined structures for different transmission electron tomography methods: resin embedded STEM-tomography and cryo-TEM tomography. The method is freely available in TomoJ software.

## 1 Introduction

Electron tomography is a technique that produces structural information of samples at a resolution between 20Å and 5Å (Kovtun et al., 2018; Mattei et al., 2018). This structural information helps scientists to determine the precise organization and relationship between the different components of samples. In biology, it has helped in understanding subcellular organelles thereby becoming a cutting-edge tool in structural biology (Donohoe et al., 2006; Pfeffer and Mahamid, 2018). In material science, the use of electron tomography is more recent, but it has already facilitated the studies going from carbon nanotubes doped with nitrogen (Florea et al., 2012) to dislocations in minerals or semiconductors (Midgley and Dunin-Borkowski, 2009).

Electron tomography relies on the tomographic reconstruction of a sample from projection images acquired at different orientations by using a transmission electron microscope. The most popular and simplest data collection geometry is the single-axis tilt series, although other geometries are also possible like dual-axis (Mastronarde, 1997), multiple-axes (Messaoudi et al., 2006; Hata et al., 2011), or conical tilt (Zampighi et al., 2005). Since single-axis tilt series is the most common and simplest method, in this paper we will focus on this technique. In single-axis tilt series, the sample is tilted around an axis to provide its different projection views. This simple approach presents some inconveniences such as the limitation of tilt range, due to the increment of the sample section to be crossed by the electrons at high tilt; mechanical instabilities during data collection producing shifted images; or deformations of images, due to small magnification change or radiation damage. Because the reconstruction algorithms are based on the existence of a single and well defined axis, the relative projection directions for each image must be established before starting the three-dimensional reconstruction process. This calls for the determination of the common tilt axis on all images, which requires the correction of shifts and/or deformations occurring during the acquisition.

Software for acquisition of single-axis tilt series is improving concerning the tracking of the sample during tilt series acquisition. However, it does not absolutely compensate for all shifts in the sample. It also tracks and corrects for changes in the defocus, but cannot compensate for magnification changes or deformations occurring in samples sensible to radiation damage. So post-acquisition alignment is mandatory to get images in the correct geometric system for reconstruction.

With biological samples, alignment of the tilt series is often performed by using gold beads serving as fiducial markers. This technique is quite effective, but since markers are not always visible or trackable, or because they are not available, it is not always applicable. In addition, the use of markers needs the intervention of user, during image processing, to manually indicate the position of fiducials on images making possible automatic tracking on the whole data set. Some papers proposed automatic detection of fiducials (Amat et al., 2008; Mastronarde and Held, 2017; Han et al., 2018). Alternatively, image corners can be used as fiducial markers (Brandt et al., 2001), or the image information content itself can be used through crosscorrelation (Frank and McEwen, 1992). Electron microscopy of single particles has proved that algorithms based on image information such as cross-correlation provide the best results when a reference volume pre-exists. However, since the objects to be reconstructed in electron tomography are unique, reference volumes cannot be generated. Consequently, post-acquisition alignment requires algorithms based on information exclusively contained in images from the tilt series. Three main possibilities using cross-correlation to align tilt series exist. The first one is to align the first image with the second, the second with the third, and so on. This serial approach compares similar images, although it has the drawback of serious potential drifts due to error propagation as it reduces the alignment to multiple two-images problems, instead of considering the tilt series as a whole. This results in inaccurate estimations of the tilt axis and shift parameters for each images. The second approach computes a rough reconstruction of the tomogram and realigns the tilt series with respect to reprojections to iteratively refine the computed reconstruction. This approach, frequently used in single particle analysis (Frank, 2006), is highly time consuming and limited for tomography because of the small number of projections available to compute the first volume. The third option is to use cross-correlation to track features within the tilt series. Features are characteristic points in the images (due to change of contrast, borders, corners, etc.) that can be tracked in neighboring images. The feature tracking allows defining markers that are visible in sub-sequences of the whole tilt series.

In 2009, we proposed an approach, based on the third option, whose improvements are discussed hereafter (Sorzano et al., 2009). This approach was as follows. First, shifts were roughly corrected by cross-correlation. Rotations and deformations were estimated using affine transforms. This was a pre-alignment required for further steps. Second, a grid was applied on each image to place seeds in regular positions. As the grid was applied on unaligned images, the seeds were not located on the same features. Third, a patch was extracted and, using the pre-alignment parameters, the corresponding patch in previous and next images was found by cross-correlation. At this step, while the correlation score between patches was high enough, the process continued to produce a landmark chain. Fourth, for each seed only chains having a length higher than a threshold defined by user were selected. Fifth, a 3D model of landmarks was created iteratively along with detection of tilt axis orientation, in-plane shifts and rotations (incorrectly detected landmark chains were automatically identified and removed).

In this article we introduce several improvements to the previously described procedure:

1. We have speeded-up the prealignment of micrographs by a factor 100 using a multiscale image alignment algorithm (see Secs. 2.1.1 and 3.2.1).
2. We have enhanced the generation of seeds producing fewer and better seeds (see Secs. 2.1.2 and 3.2.2).
3. We have introduced an algorithm specially suited to spherical image features (like gold beads, but not limited to these; see Secs. 2.1.2 and 3.2.3).
4. We have made the landmark chain generation more robust by considering the reversibility of the landmark matching (see Sec. 2.1.3).
5. We have improved the fusion of landmark chains producing longer chains and of better quality (see Secs. 2.1.4 and 3.2.4).
6. We have extended the deformation model to include sample shrinkage and shearing effects as well as differences in image magnification (see Sec. 2.2.1).
7. We have included a better procedure to spot incorrectly identified projections of the 3D landmarks or incorrect parts of the landmark chains (see Sec. 2.2.2).
8. We have modified the 3D reconstruction algorithm to account for the sample out-of-plane deformations. Alternatively, we also provide two faster 2D transformations of the experimental images that allow partial correction of the out-of-plane effects. The three alternatives are shown in Sec. 2.3, and their results in Secs. 3.1 and 3.2.5.

We finally show the performance of the method on CryoEM and resin-embedded STEM data.

## 2 Methods

In this section, the improvements for each step of the workflow of the algorithm published in Sorzano et al. (2009) are detailed. They have been included in the ImageJ (Abramoff et al., 2004; Schneider et al., 2012) plugin TomoJ (Messaoudi et al., 2007) and are freely available from http://www.cmib.fr/en/download/softwares/TomoJ.html These modifications affect the method used to generate landmark chains (Section 2.1), the deformation model (Section 2.2), and the 3D reconstruction algorithm that compensates for these deformations (Section 2.3).

### 2.1 Feature tracking to produce landmarks chains

The first part that have been improved is the one producing landmarks chains from the tilt series. It can be decomposed in four steps: 1) pre-alignment to help future tracking, 2) detection of points of interest, 3) creation of landmark chains, and 4) selection of high quality chains.

#### 2.1.1 Pre-alignment procedure simplification

In order to reduce the computation time, the entire procedure, including affine transform, can often be replaced by a simplified two-steps multiscale translation determination by cross-correlation. In this new procedure, full size images are downsampled by a factor 2 leading to an improvement of the Signal-to-Noise Ratio (SNR) as expected for an averaging filter. Then, an alignment by cross-correlation is serially performed. In addition, because the dimension of images is reduced, this leads to a gain in the computation time. The user can chose to repeat this step or to change the downsampling factor, but, as this step aims to correct for the largest translations, it is not really required. Following this first step, refinement is done by computing the cross-correlation on the center of the images at full resolution (without downsampling). Using only this central part prevents inaccuracies due to the apparition, at high tilt angle, of features in the field of view which are absent on images recorded at angles close to 0°. Sec. 3.2.1 shows the results obtained by this improvement.

#### 2.1.2 Seeds detection

Our algorithm introduced in 2009 was based on an initialization process in which the seeds were regularly placed on the intersections of a grid whose size was defined by user. However, this does not consider any information coming from images and, as a consequence many seeds were located on places with no feature to be tracked reducing the efficiency of the next steps of the workflow. Therefore, in order to optimize the initialization process, the local extrema (minima or maxima depending on image contrast) are now used as seeds. Because these extrema correspond to regions having high contrast, such as recognizable features in images or gold beads frequently used as fiducial markers, this provides an additional value to the efficiency of the determination of chains. The detailed procedure implemented consists in band-pass filtering images to remove noise, followed by the generation of a coordinates map with the position of all local minima (or maxima). This map is computed by determining if each pixel value corresponds to a local minimum within a radius given by user. For each minimum, its score is computed as the mean square difference between this pixel and its neighbors within the radius previously used. The best *N* seeds are kept, being *N* a number selected by the user. Sec. 2.1.2 shows the results obtained by this algorithmic improvement.

In addition, an optimized procedure for spherical feature detection is available. As spherical 3D features will approximately project on images as 2D Gaussians, for each detected minimum, two 1D Gaussian curves are fitted to the sum of pixel values on X and Y directions respectively. This fit is done over the radius used to determine the minimum. The centers of the Gaussian curves help determining the true center of the feature. Based on the regression coefficient used as a fit score, this approach can also be used to remove non-spherical features using a score threshold defined by the user. This optimization in the selection of spherical features directly leads to a gain in computation time for the tracking part by removing uninteresting seeds (see Sec. 3.2.3).

#### 2.1.3 Generation of landmark chains

The robust determination of landmark chains is a major issue to guarantee the final accuracy of the alignment process. In our previous work, the seeds were followed by using local patches and cross-correlation and multiple refinements of the chains, in forward and backward direction, were used to validate the chains. However, as proposed by Castaño Díez et al. (2010), this can be done in a more efficient way by checking if the matching of each landmark is reversible. This means that: given a patch located in position **p**_*i*_, in image *i*, and its corresponding matching position **p**_*i*+1_, in image *i* + 1, the matching position of **p**_*i*+1_ in image *i*, 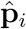 should be at the same position than **p**_*i*_. Ideally the distance 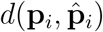 should be zero. However, this ideal case is unreachable in a large number of cases, but moderate values still gives usable matches. A maximum distance of 2 pixels is considered good enough to validate the matching as correct. Since the number of cross-correlations needed in this process is lower than that required for the procedure proposed in 2009, this procedure results in a reduction of the computation time.

In case of spherical features, an additional optimization of the landmark chains generation is possible by applying symmetry to the patch used as reference during tracking. This helps the correct tracking of spherical features when other highly contrasted structures (such as cell membranes in stained biological samples) appear in a same patch by forcing a circular shape without need of *a priori* knowledge about its size. This symmetrization is done by averaging 12 rotated images on 360° with 30° steps.

#### 2.1.4 Selection of high-quality chains

The correlation score between patches used during the generation of landmark chains is highly dependent on the matching of the patches, but also on the surface occupied by features in the patch and the SNR of the images. This score is used as the criterion to stop the tracking process when it is lower than a threshold defined by the user. However, because of the variable contribution of the three enumerated factors, it is difficult for any user to propose a correct threshold *a priori*. This difficulty results in an unpredictable computation time and in a number of chains highly dependent on the user input. In order to make the process more robust and reproducible, the user threshold is replaced by the length of the landmark chains. Thus, all seeds are tracked on a given number of images, corresponding to the selected length of the landmark chains, even if the correlation score is low. Each chain receives a score corresponding to the lowest correlation value obtained during tracking. Only chains with the highest scores are kept. The user defines the number of chains to keep, with the possibility to fix a minimum correlation value. This minimum correlation value guarantees that chains with very low correlation values, corresponding to inaccurate tracking, will not deteriorate the convergence of the alignment in the next step.

In addition, as some of the local minima can fall on a same structure in the sample, many landmarks chains can share a common segment, which leads to an over-representation of some regions of the images. To prevent this, a fusion of landmark chains is done when the landmarks chains share landmarks within a radius of less than two pixels on at least three images. The fusion is done by averaging the position of landmarks. This procedure has the advantage to produce longer chains than those defined by user, allowing a significant reduction in the number of images on which seeds might be tracked. This new approach makes, in a robust way, the selection of chains more independent of arbitrary and difficult to define parameters. Sec. 3.2.4 shows the results related to this part of the algorithm.

### 2.2 3D Deformation model and estimation of its parameters

The improved landmark chains are then introduced into an alignment algorithm that will create a 3D model with the 3D coordinates of each landmarks. This 3D model was improved by adding sample deformations, and variable magnifications.

#### 2.2.1 Alignment with normalized distances

Let **r**_*j*_ be the coordinate of a 3D landmark, **p**_*ij*_ the coordinate of its projection onto image *i*. Let *V*_*j*_ be the set of all images in which the *j*-th landmark is seen, and *V*_*i*_ the set of all landmarks seen in image *i*. The relationship between **r**_*j*_ and **p**_*ij*_ is

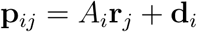

where *A*_*i*_ is the projection matrix accounting for the tilting around the tilt axis and a posterior in-plane rotation, and **d**_*i*_ is a 2D vector accounting for an in-plane shift of the *i*-th image. *A*_*i*_ is computed as

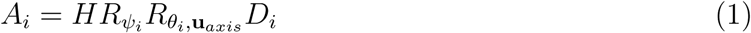

where *H* is the matrix that projects 3D coordinates into its *X, Y* components, 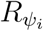 is a rotation matrix of *ψ*_*i*_ degrees around the *Z* axis (that is, the beam axis), *D*_*i*_ is any 3D deformation matrix with which we can encode change of scale (with a diagonal matrix) or shearing along any arbitrary axis (a unitary matrix with all its eigenvalues equal to 1 and a degenerate eigenspace) or any other arbitrary 3D transformation, and 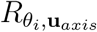 is a rotation matrix of *θ*_*i*_ degrees around the tilt axis **u**_*axis*_. The tilt axis is described by the two Euler angles *α* and *β* yielding the vector representation

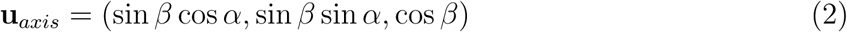

Let us also refer to the product 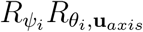 as *R*_*i*_. In the Electron Tomography field, it is traditional to model *S*_*i*_ (Mastronarde, 2006) as

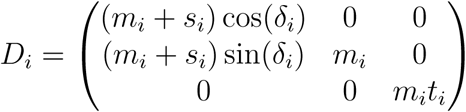

although we prefer the model

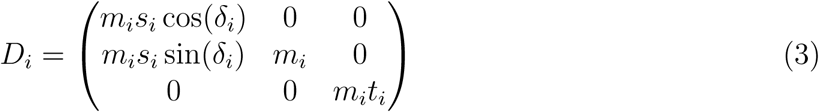

Both deformation models can span the same subspace of deformation matrices. In this model, *m*_*i*_ is a global change of magnification/compression, *t*_*i*_ is a vertical compression of the specimen, *s*_*i*_ is a compression along *X*, and *δ*_*i*_ represents a shearing operation.

The goal is to minimize the reprojection error

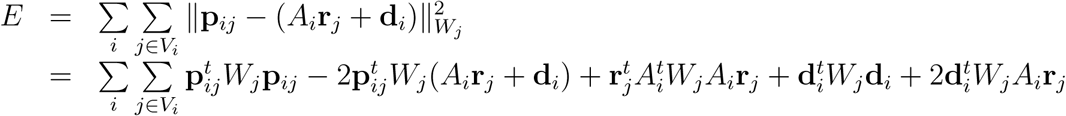

where 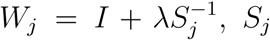 is the sample covariance of the residuals of the *j*-th landmark calculated as

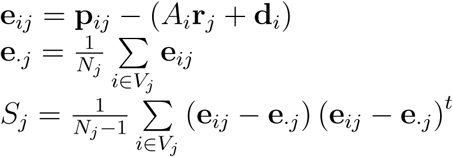

and *λ* is a regularization parameter that balances the importance between the Euclidean and Mahalanobis distance (we have observed that a weight different from zero helps to estabilize the alignment, especially by avoiding jumps in the zero degree image). In fact, this goal function is a generalization of our previous one, since the previous corresponds to the particular case *λ* = 0. Note that *W*_*j*_ is a symmetric matrix 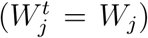, and that at every iteration of our algorithm we only have an estimate of the matrix 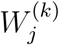 given by the current estimates of the parameter 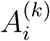 and 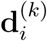.

In order to reduce the cross-talking between alignment parameters, we constrain the minimization so that

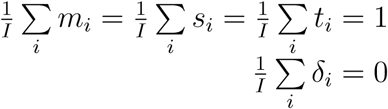

which is simply stating that the mean of each of the deformation parameters should be around 1 (no scaling, *m*_*i*_, *t*_*i*_, *s*_*i*_) or 0 (no shearing, *δ*_*i*_).

For performing the minimization of *E* with respect to the different parameters, we differentiate *E* with respect to each parameter and equate to 0. This gives us updates for all the model parameters (different deformations of the volume, image shifts and rotations, 3D landmarks, and position of the tilt axis). The details of these updates can be seen in the Suppl. Material 1.

#### 2.2.2 Evaluation of the landmarks quality and removal of outliers

We can also use the normalized distance introduced in this paper to evaluate the quality of the landmarks used for alignment. For doing so, we can analyze the residuals, **e**_*ij*_, which we have seen that play a central role along the alignment procedure, and identify those landmarks with larger residuals according the metric introduced in this paper. We can do this in two different ways:

1. <u>Identification of large isolated residuals</u>: We may identify those landmarks for which at least one of their residuals in at least one of the projections is larger than a given threshold:

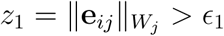
2. <u>Identification of large average residuals</u>: We may identify those landmarks for which, on average, their residuals are larger than a given threshold:

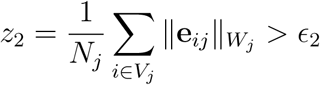

These thresholds may be either set by the user or estimated by some suitable procedure like calculating a high percentile (90-95%) of the distribution of the measured feature (‖**e**_*ij*_‖*W*_*j*_ or 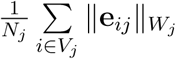) over all landmarks. Alternatively, we here propose an automatic way of removing landmarks based on Mahalanobis distance. Let us define for landmark *i* a vector formed by its residuals 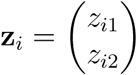. We define the covariance of all residual vectors as

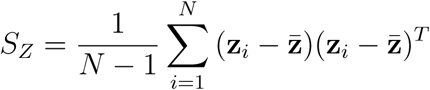

where *N* is the number of landmarks and 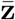 is the mean of all residuals. Then, for every landmark we compute its Mahalanbobis norm with respect to the distribution mean as

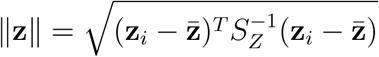

Those landmarks whose norm is larger than a threshold are identified as outliers. In the multivariate statistics literature it is normally accepted to use the threshold 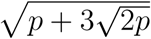, where *p* is the number of variables of the **z** vectors (in our case, *p* = 2).

### 2.3 3D Reconstruction and compensation of the deformations

Once the alignment has been estimated, we want to produce a reconstruction that takes into account the deformation in such a way that the reconstructed volume is undeformed. In the following we show three different methods with increasing degree of accuracy.

#### 2.3.1 Method 1: Linear mapping of the acquired images

Following the classical approach in the spirit of Mastronarde (2006), we may try to find a linear transformation that converts points in the underformed projection into points in the deformed (acquired) one. Consider a point **r′**in the underformed volume *V*^*′*^. In the undeformed projection it would be projected at the coordinate

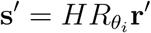

while in the deformed projection it would be projected at the coordinate

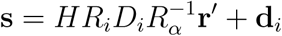

We may now look for an affine transformation of one coordinate into the other

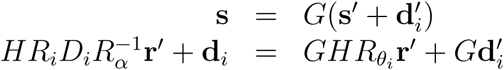

which implies

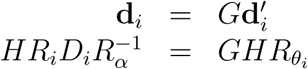

If we multiply the second equation by *H*^*t*^ from the right

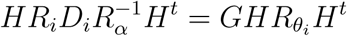

From this last equation we can solve for *G*

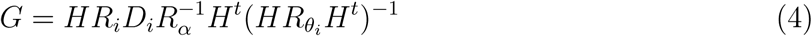

You may compare this latter equation with Eq. 5. It can be seen that the latter is a 2D version of the one in Method 2. Once we have the matrix *G* we can easily calculate the affine displacement

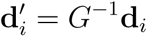

#### 2.3.2 Method 2: Non-linear mapping of the acquired images

For reconstruction, it is convenient to have the tilt axis aligned with the vertical axis and with all shifts and deformations corrected. For this, we need to generate the images

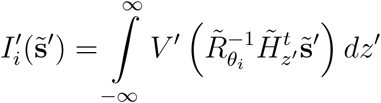

where we have made used of the matrix

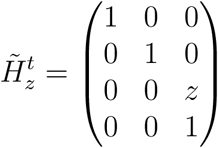

and used homogeneous coordinates (with tilde) to refer to all positions in space (as a reminder, homogeneous coordinates for a 2D or 3D location are not unique, but it suffices to think of them as 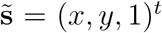 or 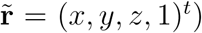. Among all 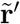 points projecting onto a given coordinate 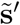 only one of them lies in the *XY* plane, 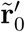

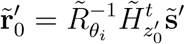

Let 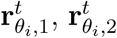 and 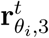 be the first, second and third rows of the 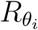 matrix. Then, it can easily be shown that (note that we have abandoned the homogeneous coordinates)

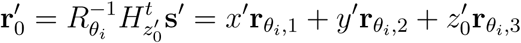

Since 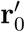 lies in the XY plane, it must be

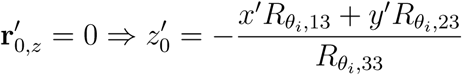

In the experimental (deformed) image, this coordinate is projected at the location

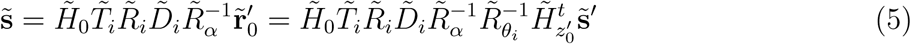

Note that this relationship is valid only for the **r**^**′**^points in the *XY* plane and that the relationship is non-linear because the matrix 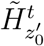 depends on the coordinate **s**′.

We can relate the ideal images needed for reconstruction with the acquired images as

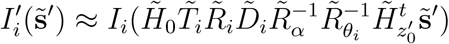

Note that because of the 3D deformation *D*_*i*_, this latter equation gives only an approximation to the true image 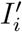. The larger the deformation and the sample thickness, the worse the approximation is. However, the approximation becomes exact for infinitely thin samples. Instead of creating “undeformed” images, a more exact alternative would be to construct a 3D reconstruction algorithm whose projector explicitly takes into account the deformation.

In the following paragraph we show that this way of generating an “undeformed” projection generalizes the current approach to projection alignment of a tilt series. In case, there is no deformation (the assumption of the standard tomographic approach), the expression above simplifies to

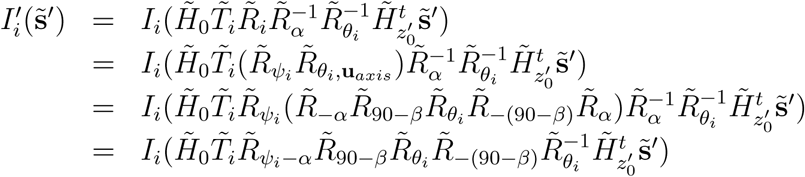

where *α* and *β* are the two angles defining the tilt axis (Eq. (2)), 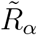 is a rotation around 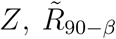 is a rotation around *X* and 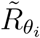 is a rotation around *Y*. If the tilt axis is perfectly perpendicular to the electron beam, then *β* = 90 and the expression above further simplifies to

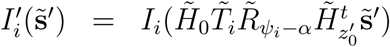

Note that the translation and rotations are only performed in the *XY* plane, so using the 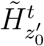 or 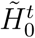 gives the same result. Consequently,

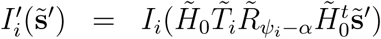

That is the standard way of aligning projection images of a tilt series.

#### 2.3.3 Method 3: 3D Projector integrated in the 3D reconstruction algorithm

Each one of the images acquired responds to a model that can be described as (Sorzano et al., 2014, 2015)

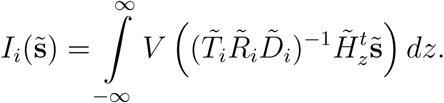

For the sake of clarity let us specify each one of the matrices involved:

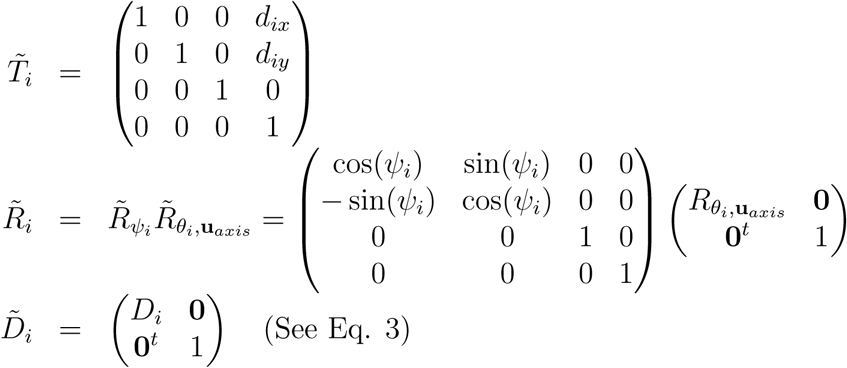

We are interested in producing a tilt series whose tilt axis is not arbitrary but it is aligned with the *Y* axis. Let us call *V′* the corresponding volume. The relationship between *V′* and *V* is given by

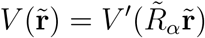

where *α* is the rotation angle defining the orientation of the tilt axis orientation (see Section 2.2.1) and 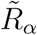 is a *Z*-rotation matrix with this angle. In our convention, angles are measured from the *Y* axis (i.e., the *Y* axis has *α* = 0) and clockwise angles are positive.

Thus, we may rewrite the acquired image in terms of the vertically aligned volume as

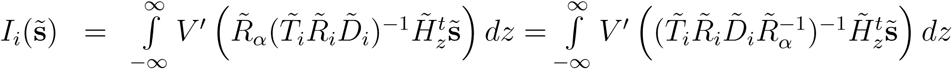

Note that all points in the volume *V′* of the form

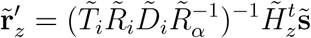

project onto the point 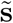 of the experimentally observed image. Conversely, given a point 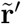 of the volume *V*′, it is projected on the experimental image at the location

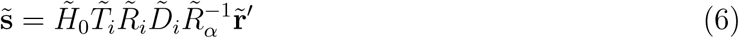

Including Eq. 6 into the reconstruction process breaks the possibility to optimize parallel beam reconstruction by working on each XZ plane independently. This drawback is particularly crucial with optimization on GPU where memory is often not sufficient to hold the entire reconstruction. The reconstruction of part of the reconstruction and then joining them is problematic in the junctions, because some of the information to use in the process is not attainable. To solve this problem, it is possible to reconstruct overlapping sub-reconstruction and then copy only the parts where all information is present. Using Eq. 6 we can determine the change for volume coordinates of each point as 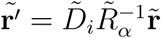

Knowing this, it is then easy to determine the range of the correctly reconstructed pixels from the corners coordinates of the sub-reconstruction.

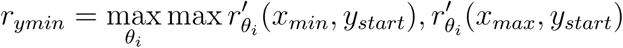

and

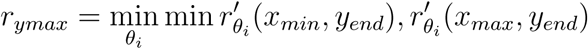

Finally, to determine the start of next sub-reconstruction, we use the same approach by taking

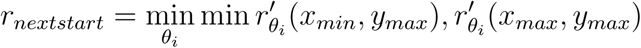

Special care with these limits should be taken when *α* is higher than 45° or lower than −45°, and in these cases we must use *α ±* 90°, in order to have the correct image corners.

## 3 Results

We first show the ability of the algorithm to recover deformed tilt series with a phantom test. This test will also help to understand the artifacts induced by the different correction methods. We then illustrate each one of the new improvements on a standard dataset (Experimental example 1) widely used in ours and others’ publications. Finally, we show the application of our method to the alignment of datasets in the absence of fiducial markers (Experimental examples 2 and 3).

### 3.1 Phantom test

Phantom data was used to validate the new model used for alignment and prove its correctness. First, a phantom volume was created as white dots randomly distributed in a dark background. The projection of the points were computed using Eq. 1 with variations of shifts, rotation, and deformation parameters following sine functions. The maximum values were 50 pixels for shift, 2° for in-plane rotation (*ψ*_*i*_), 0.02 for magnification (*m*_*i*_), 0.05 for shrinkage (*t*_*i*_), 0.1 for scale on X axis (*s*_*i*_) and 0.5° for shearing angle (*δ*_*i*_). The reader is referred to the Sec. 2.2.1 for a detailed explanation of these parameters. The coordinates on the projection images were stored and then used as input to the procedure to determine all parameters for alignment and deformation. The parameters of deformation found were very similar to the true ones (Supplementary Material 2). However, to quantitatively assess the effect of the small differences in the deformation parameters, we calculated the warping index between the true transformations applied to the phantom at tilt *i, D*_*i*_, and the estimated ones, 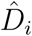, (Sorzano et al., 2005)

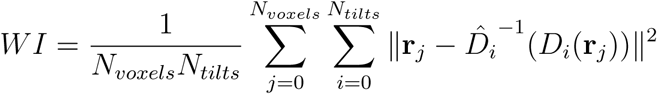

where *N*_*voxels*_ and *N*_*tilts*_ are the number of voxels in the phantom and the number of tilts images created respectively. **r**_*j*_ is the *j*^*th*^ voxel coordinate in the phantoms. On the phantom data used in Fig. 1 the warping index was 0.24 voxels. This difference is in the subvoxel accuracy, proving the accuracy of the algorithm.

**Figure 1:**
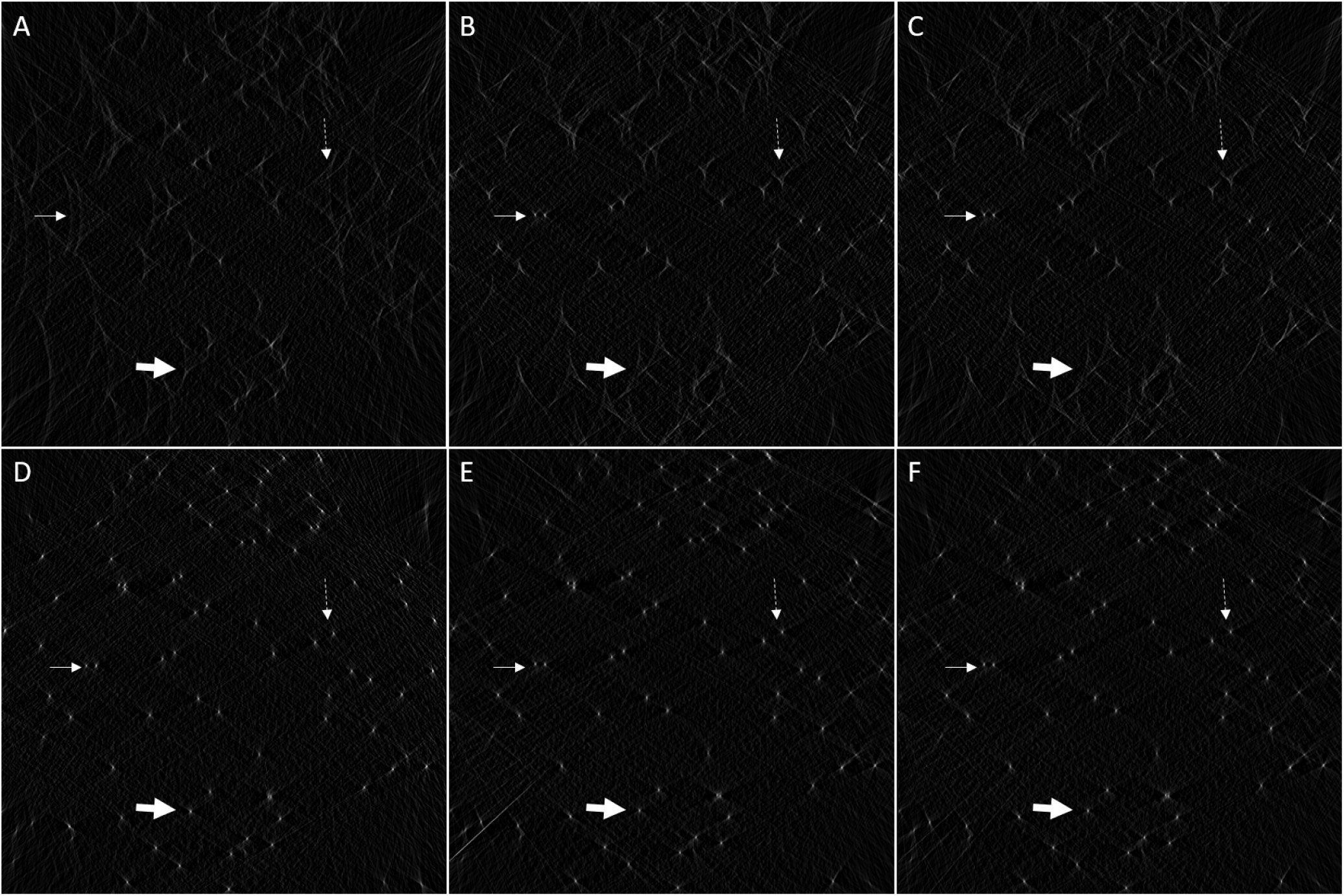
Comparison of the reconstructions of the phantom with the different method of correction of deformation. Each image corresponds to the sum of XZ planes to better see the artifacts: (A) no correction of deformations, (B) correction using 2D affine transforms, (C) correction using 2D non-linear mapping, (D) optimal reconstruction if no deformation exists, (E) correction using 3D projector integrated in reconstruction, (F) correction using 3D projector integrated in reconstruction with deformations computed from landmarks. White arrows point to features. Near the central plane (thin arrows), the feature are well reconstructed with all methods of correction of deformation. The features above (dotted arrows) or far (large arrows) from central plane display correction of the in-plane deformation, but show triangular shapes unless the 3D projector is employed.

In Sec. 2.3, we discuss three different ways of correcting the estimated deformations. To show their effect, three different reconstructions were performed using the true deformations parameters. As can be seen in Fig. 1, the points on the central plane are perfectly reconstructed with the different methods of correction of deformations (thin white arrows). The points that are not on the central plane are perfectly reconstructed only in the case of applying alignment with the 3D projector during the reconstruction process (Method 3; Fig. 1 E and F). With the other two methods (B and C), the points are reconstructed as triangles (dotted or large white arrows).

### 3.2 Experimental example 1

In Sec. 2.1 we have introduced several modifications for the detection and tracking of landmarks. In this section we test these ideas along with the analysis of the different ways of estimating (Sec. 2.2) and correcting (Sec. 2.3) for the deformation. To evaluate these ideas, several tests were performed on the TomoJ tutorial data (*Pyrodictium abyssii*) cell strain TAG11 (Rieger et al., 1995), embedded in vitreous ice layer on holey carbon grid (Sorzano et al., 2009).

#### 3.2.1 Multiscale prealignment with cross-correlation

The total computing time was reduced from 388 s. in our previous method to 4 s. in the method introduced in this paper using a laptop computer equipped with an Intel i7-4700MQ processor and 32 GB RAM. This implies a speed-up factor of two orders of magnitude. We measured the quality of the new prealignment by evaluating the number of landmark chains and their alignment score. The alignment score of the previous algorithm (Sorzano et al., 2009) and the new one are very similar, 1.37 and 1.38, respectively. The alignment score is the average error in pixels between the reprojection of the 3D landmarks and their experimentally observed positions. However, the number of landmark chains dropped from 2,684 to 2,127. Although there is a drop of about 20% in the number of chains, there is no drop in the quality of the detected chains, and the execution time has been reduced by a factor 100.

#### 3.2.2 Seed generation

We evaluated the ability of the new methods to generate seeds using the ratio of final retained chains versus the number of initial chains (Table 1). We see that in our previous method the success rate of a seed was about 5.8% that was the probability of a grid point hitting a feature that could be tracked along the tilt series. We see that the new method is much more successful resulting in a larger number of final chains and higher success probability. The fact that adding more seeds per image in the new method reduces the success rate can be explained by the fact that high quality seeds are chosen first, and adding more seeds per image results in a degradation of the quality of the added seeds.

**Table 1:**
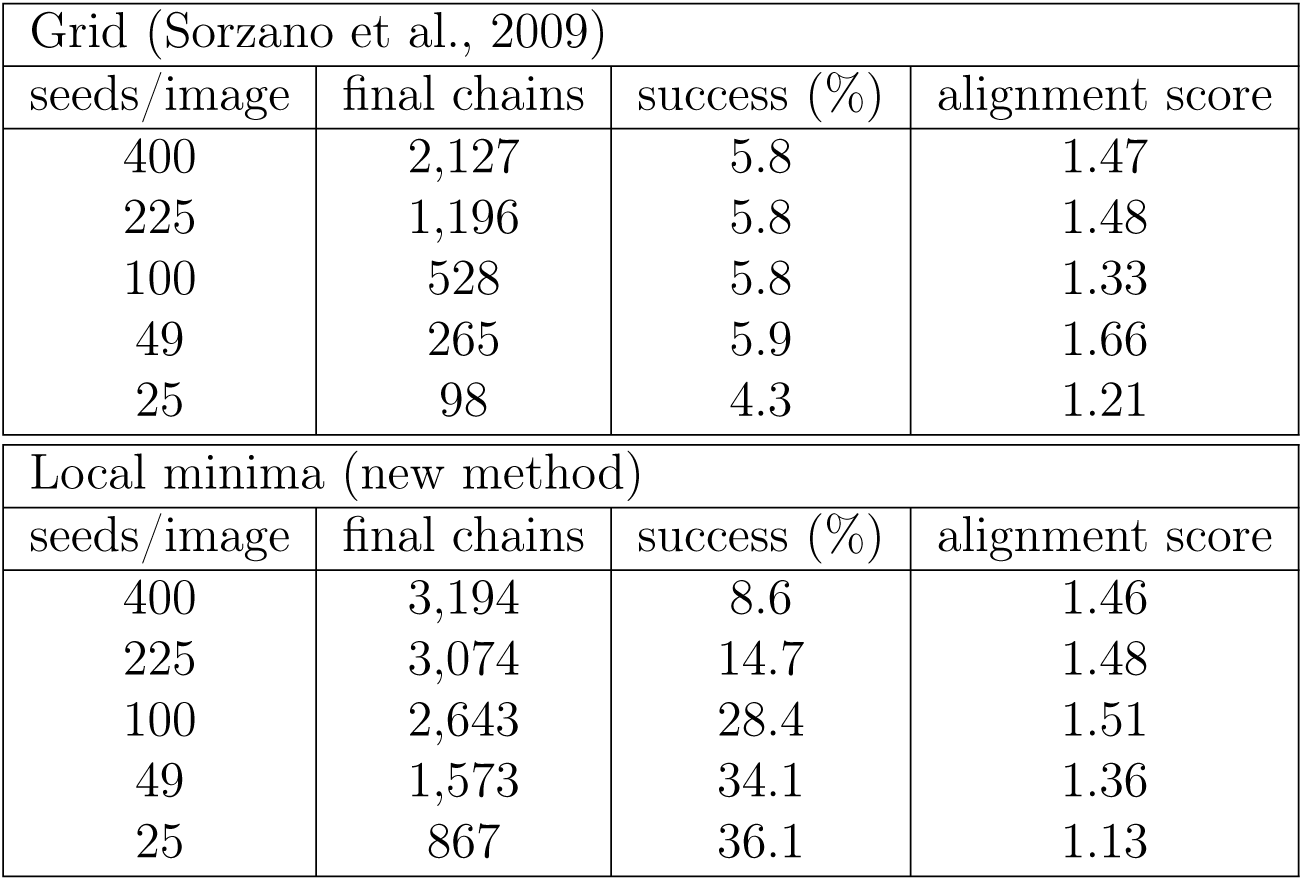
Comparison of the method of seeds determination. For our previous and current methods. The table shows the number of seeds per image, the number of final chains, the success probability of a seed to generate a final chain, and the alignment score in pixels.

#### 3.2.3 Spherical feature optimization

The spherical feature optimization plays a role at two levels. First, the detection of seeds by filtering with Gaussian function. The effect of this step is to reduce the number of seeds to track in the subsequent algorithm. The activation of this option with a threshold of fit of 0.5 gives 5,792 seeds instead of the 18,592 found after the detection of local minima. The number of chains found dropped from 7,708 to 3,176 (1,638 and 943 after fusion respectively) and the final alignment score is very similar: 1.26 and 1.20 without and with gold bead optimization, respectively. The main gain is the computation of only 31% of the potential seeds at the expense of computing the Gaussian fitting. This is confirmed by the total time to compute alignment (detection and alignment) that drops from 10 minutes and 22 seconds to 4 minutes and 46 seconds.

The second role of spherical feature optimization is in the tracking of the seeds. The effect of the symmetrization during tracking resulted in a lower number of chains found, 1,992 (166 after fusion), compared to the tracking without symmetrization, 3,176 (943 after fusion). First, the fusion is more efficient with the symmetrization with only 8% of the chains remaining instead of 29%. Second, after fusion the length of landmark chains improves from an average length of 35 to an average length of 47. The number of tracked chains highlights the fact that the symmetrization limits the selected features to spherical ones. The high level of fusion proves that few distinct features are found by the procedure. Moreover, the interest of the procedure for spherical features is confirmed by the increase of average length of landmark chains as long chains help the future alignment. Adding this symmetrization step allows a new drop in the total computation time to 2 minutes and 55 seconds.

As can be observed in Fig. 2 the algorithm found many spherical features of different sizes. Some of them are gold beads present on sample as expected. However, many are just particles present in ice or even proteins of the archaeon.

**Figure 2:**
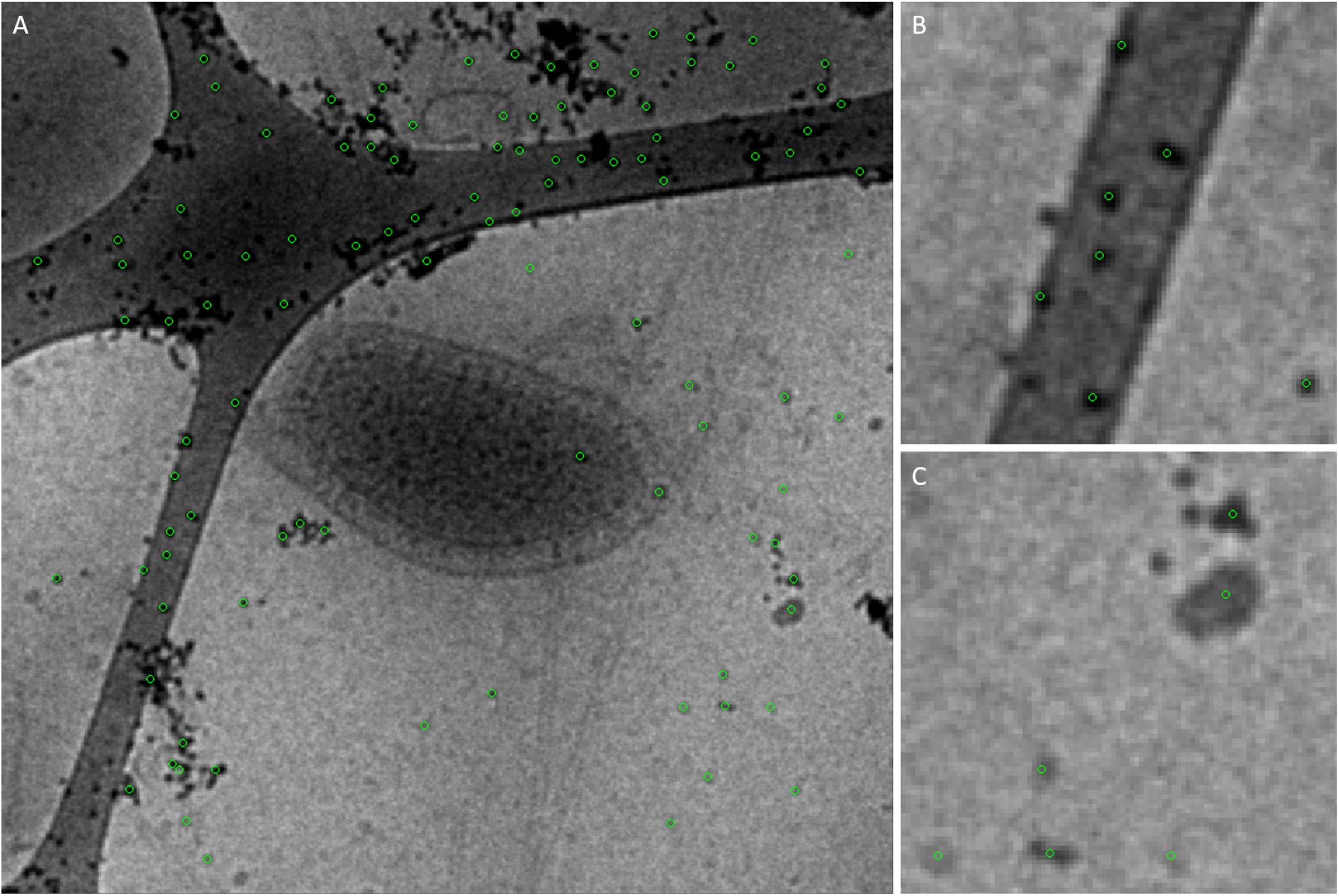
Visualization of landmarks chains found with the gold bead-like optimization. A) The landmarks found are displayed with green circles on the projection image near 0°. B) zoom on a region with gold bead. C) zoom on a region with no gold bead, but sufficiently spherically symmetric, distinctive features.

#### 3.2.4 Tracking of landmark chains

In Sec. 2.1.4, we have proposed a cross-validation procedure to track a seed along the different tilt series images. The use of this procedure increased the number of chains from 3,036 to 3,701 (increase of 21%) while keeping similar alignment score (1.48 and 1.54 respectively).

The use of fixed sized chains proves to be efficient. A first experiment was done with the correlation threshold approach with a minimum size of 15 and threshold of 0.9. It produced 3,702 chains with an average length of 52 and correlation scores ranging from 0.90 to 0.96. For the fixed size approach, chains of size 31, the algorithm produced twice the number of chains (7,708) with correlation scores ranging from 0.5 (minimum accepted score) to 0.96. After alignment, the score obtained slightly improved (1.09 for fixed size and 1.54 for the old approach).

The fusion of landmarks reduced the number of chains from 7,708 to 1,638 chains. The fused chains had an average length of 39 images. The chain fusion on the alignment caused a slight increase of the alignment score from 1.09 to 1.25.

#### 3.2.5 New reconstruction algorithm with deformation compensation

The new reconstruction algorithm was first compared with the previous version. The reconstructions, Fig. 3, show an important improvement. As can be seen, the gold beads pointed by white arrows display a higher circularity and no banana shape.

**Figure 3:**
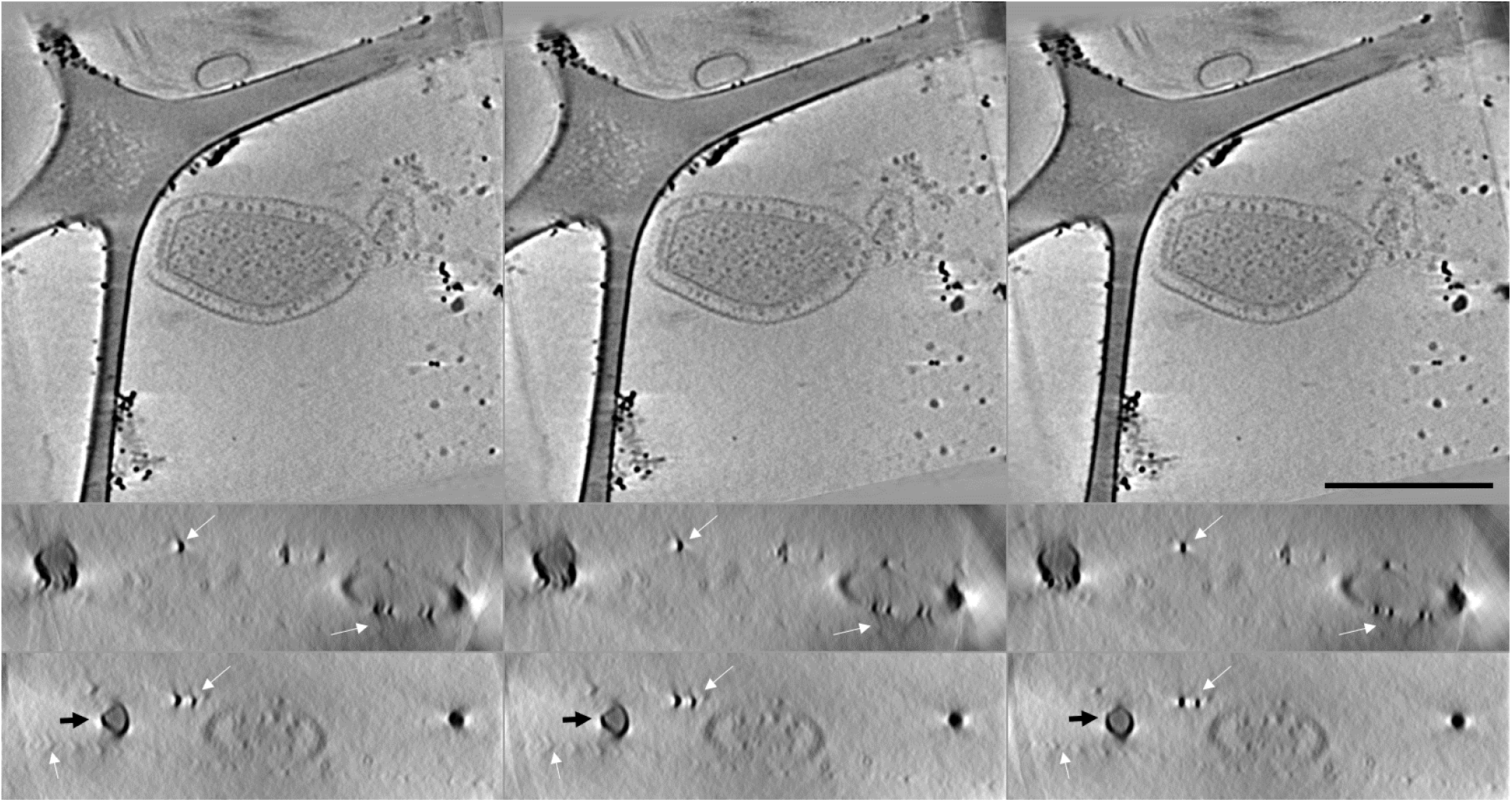
Comparison of reconstructions obtained with different alignment models. The first column corresponds to alignment using our old alignment model (Sorzano et al., 2009), the second column displays the result of the new model with only shifts and in-plane rotation (this experiment shows the improvements not due to the 3D deformation model), and third column shows the result of the new model with deformation correction. The top row corresponds to the central XY plane. The second and third rows correspond to the XZ plane number 49 and 315, respectively. White arrows points to features with visible change between algorithms. Black arrows points the plastic of holey grid where change in shape is visible. The scale bar corresponds to 500 nm.

Next, we explored the effect of the three different deformation compensation algorithms. The three compensation schemes were tested. As Fig. 4 shows, visible changes between the three methods are present. In the case of application using 3D projector during reconstruction (Method 3) the gold beads pointed by white arrows display less reconstruction artifacts.

**Figure 4:**
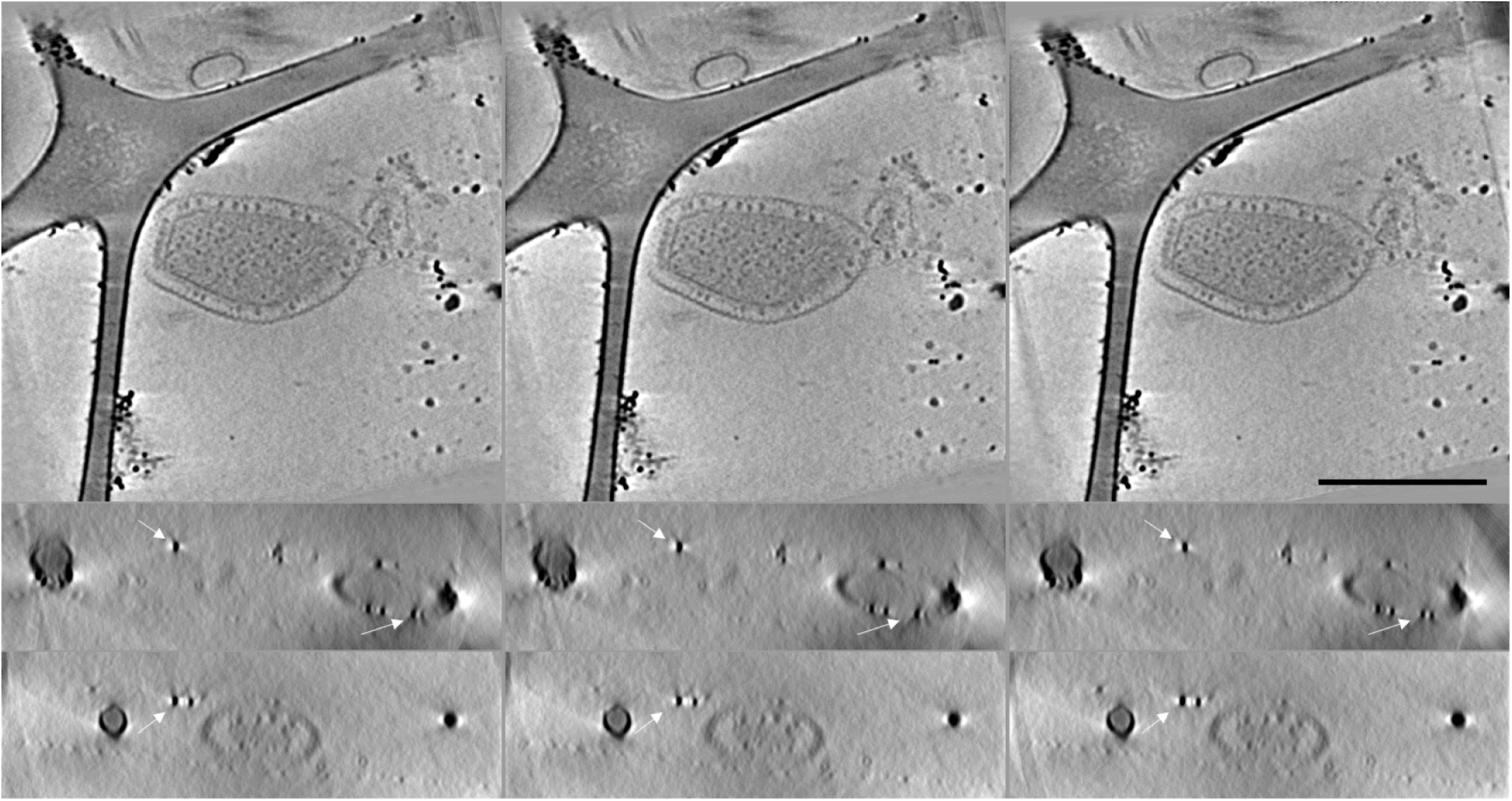
Comparison of the methods of application of alignment with deformations. The top row corresponds to the central XY plane. The second and thirds row correspond to the XZ plane number 49 and 315, respectively. The first column corresponds to method 3 with application during reconstruction process. The second column corresponds to method 2 with non-linear mapping. The third column corresponds to method 1 with linear mapping. White arrows point to features with visible change between algorithms. Scale bar corresponds to 500 nm.

### 3.3 Experimental example 2

The method was applied on an acquisition of *Trypanosoma brucei* cilia embedded in Epon resin. The electron tomography was performed using a JEM 2200FS microscope (JEOL, Tokyo, Japan) operated at 200 KeV in STEM bright field mode. The tilt series ranged from −75.62° to +77.62° with a Saxton scheme (Saxton, 1978) for a total number of 97 images. The images, of a size of 2, 048 × 2, 048 pixels, were recorded at nominal magnification of 100,000X with a pixel size of 1.46 nm. at specimen level. Images were binned by a factor 2 before processing. The algorithm was applied using the following parameters. For prealignment, the two pass cross-correlation approach was applied on a region of interest (ROI) centered on image center. First pass was with a size of 720 × 720 pixels followed by a factor 2 binning. The second pass was performed on a 360 × 360 pixels size ROI with no additional binning. The generation of landmarks was done using local minima detection (radius 4 pixels) on band-pass filtered images (with radius in Fourier images 0 and 256 pixels). 1,800 seeds were detected on each image (on the 95% central part). The tracking length was 24 images using a 11 × 11 pixels size for the patch with a minimum allowed correlation of 0.5. After tracking, 111,963 chains were found. After fusion of these chains, 24,575 remained.

Two independent alignments were produced from the fused chains. The first was with only shift and rotation, the second with all the deformations defined in Eq. 3. With only shifts and rotation the final score was 1.99 (average residuals 1.20, isolated residuals 2.79) with 18,083 chains remaining. With deformations, the final alignment score was 1.81 (average residuals 1.09, isolated residuals 2.53) with 21,243 chains remaining. Reconstructions were done using SART algorithm with 20 iterations and a relaxation coefficient of 0.1. As can be seen in Fig. 5, the two reconstructions of the cilia seems at first similar proving that the alignment procedure gives good alignment. However, close examination of the reconstructions shows clear differences such as microtubules under the membrane in the bottom left corner of the central XY plane. Without correction of deformations some part of the volume are very good while others are blurred. With correction of deformations, the reconstruction is well defined in all areas.

**Figure 5:**
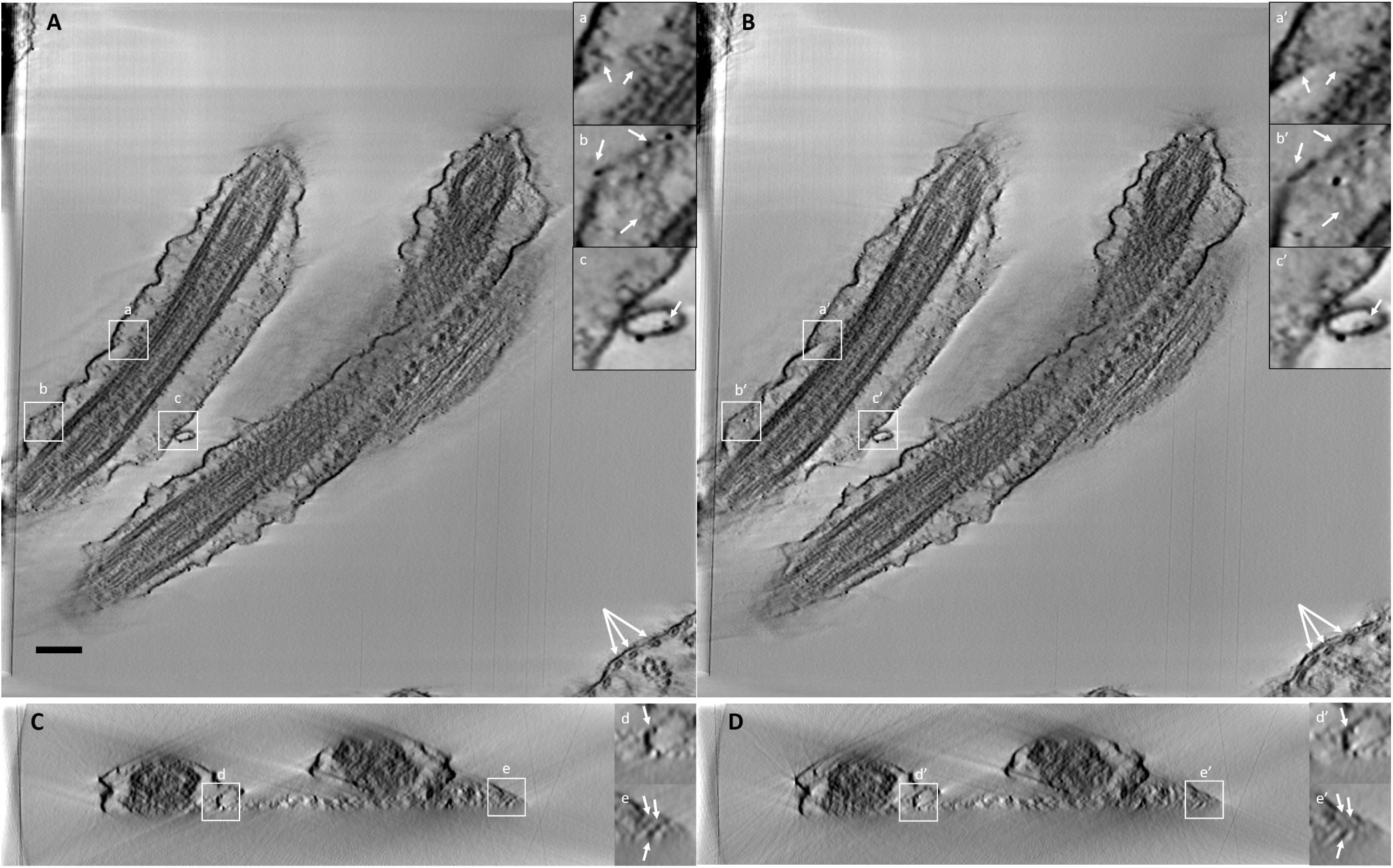
Comparison of the reconstructions of *Trypanosoma Brucei* cilia obtained with or without deformation. A and C: Central slice in XY and XZ direction respectively from the reconstruction with deformation correction, B and D: Central slice in XY and XZ direction respectively from the reconstruction without correction of deformations. a, b, c, d, e zoomed-in region corresponding to the areas delimited by white squares on central slices from reconstruction with deformation correction. a’, b’, c’, d’, e’ zoomed-in region from reconstruction without correction of deformations corresponding to the same areas as a, b, c, d, e respectively (delimited by white squares on central slices). White arrows points to details improved in the reconstruction with deformation correction. The scale bar corresponds to 200 nm.

### 3.4 Experimental example 3

The method was applied on an acquisition of T5 bacteriophages. Bacteriophages T5st(0), a heat-stable T5 mutant, were produced in E.coli Fsu*β*+ and purified on CsCl gradient as described in Boulanger et al. (1996). They were dialysed against 10 mM Tris-HCl pH 7.5, 100 mM NaCl, 1 mM MgCl2, 1 mM CaCl2, and stored at 4°C at a DNA concentration of 7 mg/ml. 3-4 *µl* of the phage suspension were deposited onto a glow-discharged cryoEM grid (Quantifoil R2/2). The grid was blotted with a filter paper for 4 s, and plunged into liquid, using a Vitrobot Mark IV operated at room temperature and 100% relative humidity. Tilt series were collected at 300 kV on a Titan Krios electron microscope equipped with Volta phase plates, a GATAN GIF Quantum post-column energy filter and K2 Summit direct electron detector, at a pixel size of 2.185 Å, using the Serial EM software. A dose-symmetric recording scheme was used, from a starting angle of 0°, and an angular increment of 2°, within a *±*60° angular range. The electron dose was set to 1.2*e/*Å^2^ for individual images, corresponding to a total dose of 73.2*e/*Å^2^. Images were binned by a factor 2 before processing. The algorithm was applied using the following parameters. For prealignment, the two pass cross-correlation was applied. First pass was with a size of 1, 024 × 1, 024 pixels followed by a factor 2 binning. The second pass was performed on a 512 × 512 pixels size ROI with no additional binning. The generation of seeds was done local minima detection (radius 15 pixels) on band-pass filtered images (with radius in Fourier images 2 and 125 pixels) with 2,000 seeds detected on each images (on the 90% central part). The tracking length was 12 images using a 41 × 41 pixels size for the patch with a minimum correlation allowed of 0.3. The algorithm identified 787 chains found after fusion. The alignment with all deformations was computed with this new chains resulting in a score of 3.33 (average residual 2.21, isolated residual 4.48) with 444 chains remaining. The reconstruction was SART with 600 iterations and a relaxation coefficient of 0.01. As can be seen in Fig. 6A, the viruses were well defined (the zoomed views are filtered using a 3D Gaussian blur of radius 1). Around half of them were intact and still contain their DNA inside (Fig. 6, B and C) while the other half were empty with broken capside. Interestingly, three of the empty viruses seemed to have some DNA chains still connected to the tail. The tails were well defined (Fig. 6, H and I) with a periodic structure clearly identifiable (Fig 6, I’) as helicoidal with a pitch of 43.6Å and a twist of 38.66°(in Effantin et al. (2006) the values described were 43.8Åand 39.4°respectively). The tail end is capped with the central fiber and lateral fibers were connected (Fig. 6, E and F). Some of the tails have lost the capping by the central fiber, they corresponded to empty viruses, even if no link to this state can be made at the moment (Fig. 6, G). The only gold bead present in this sample showed no deformation (Fig. 6, J and J’)

**Figure 6:**
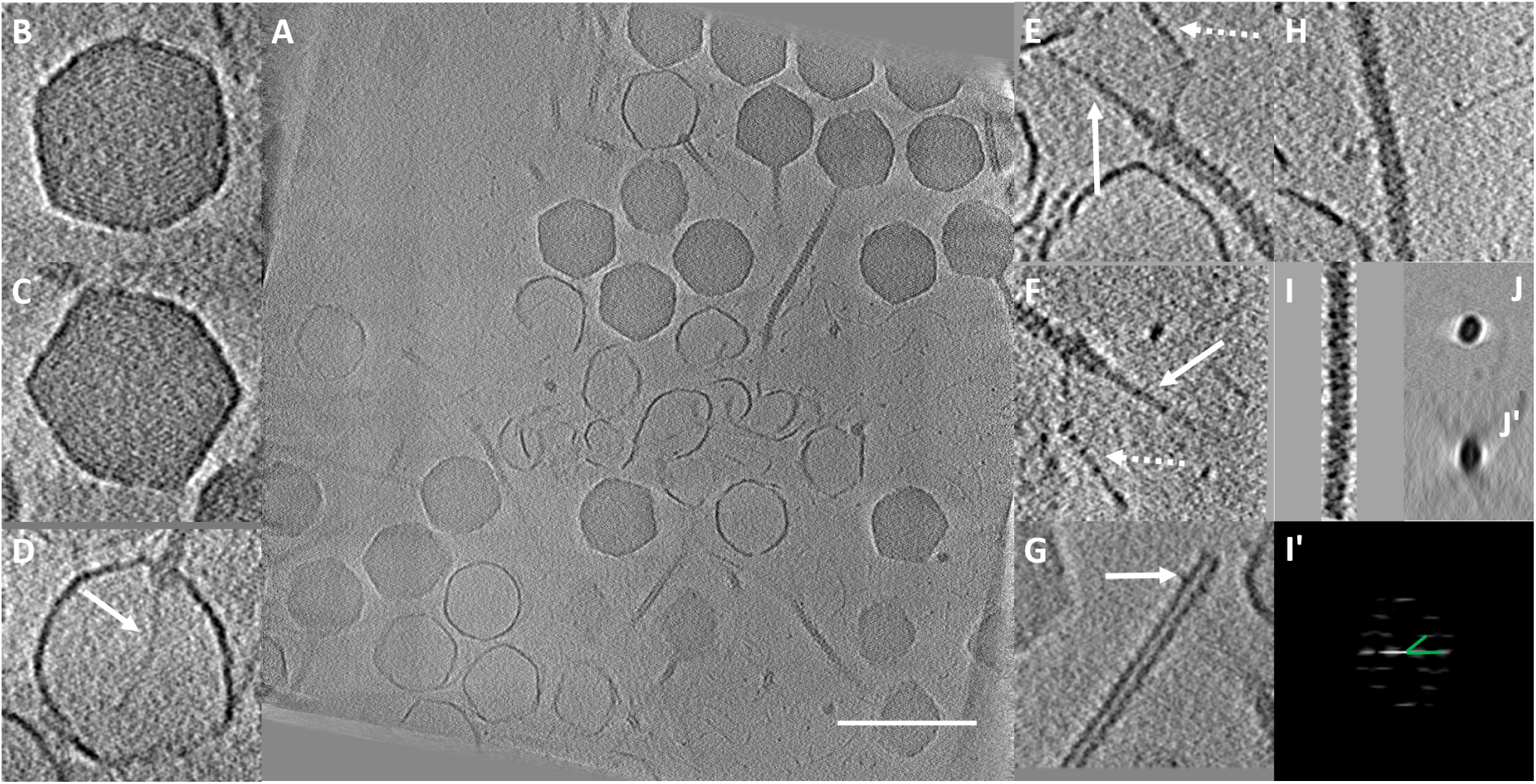
Reconstructions of bacteriophage T5. A) plane from the reconstruction. B to D zoom on virus either filled with DNA (B and C) or empty except for a brand of DNA going through the tail (white arrow). E to G zoom on tail extremity, when viruses are filled with DNA the tail fibers are clearly identifiable in E and F: central fiber is pointed by white arrow, and lateral by dotted arrow; when viruses are empty, the central fiber is lost and the tail is open. H zoom on tail, I average of a tail and I’ its Fourier transform with spots corresponding to helical structure the pitch angle (green lines) is 38°. J gold bead and j’ view in XZ plane. Scale bar corresponds to 200 nm.

## 4 Conclusions

We have shown with phantom data that the new methods are capable to properly estimate the underlying deformations. The correction of these deformations was shown to be better when done during 3D reconstruction than when applied on 2D projection images. This can be explained by the fact that the two methods correcting deformations by modifying the 2D projections consider that all points on the images come from the same plane (see Method 2 derivation). However, these two corrections provide fast approximations to the more expensive 3D full correction. The parameters used to create the phantom data are larger than the ones usually found on experimental data, thus, this artifact is much less visible in experimental data, although it is present.

The differences between the affine transform and the nonlinear transform are small, even if in theory the nonlinear version should be better. The main drawback of the affine transformation is that it is correct only in the central plane, but not in the other planes of the reconstruction. On the other side, the nonlinear correction is computationally more demanding as the step to create the aligned image is more complex.

On experimental data, the improvement of seed detection and tracking helps to produce better chains with easier parameterization of the algorithm. The alignment with deformations proved to be worthy even on cryo-tomography data. The algorithm permits a detailed analysis of the different parameters in the deformations. The correction of the deformation proved to be more efficient with Method 3. The method proved efficient in aligning cryo-tomography data without the need of gold beads.

## Acknowledgements

We thank S. Trepout for the STEM dataset on cilia of *Trypanosoma brucei*, M. de Frutos (LPS, Orsay) for phage purification and J. Ortiz (IGBMC, Strasbourg) for tilt series acquisition.

We thank financial support from the Spanish Ministry of Economy and Competitiveness through Grants BIO2016-76400-R(AEI/FEDER, UE), the “Comunidad Autónoma de Madrid” through Grant: S2017/BMD-3817, Instituto de Salud Carlos III, PT17/0009/0010 (ISCIII-SGEFI/ERDF), European Union (EU) and Horizon 2020 through grants: CORBEL (INFRADEV-1-2014-1, Proposal: 654248), INSTRUCT-ULTRA (INFRADEV-03-2016-2017, Proposal: 731005), EOSC Life (INFRAEOSC-04-2018, Proposal: 824087), HighResCells (ERC-2018-SyG, Proposal: 810057), IMpaCT (WIDESPREAD-03-2018 - Proposal: 857203), EOSC-Synergy (EINFRA-EOSC-5, Proposal: 857647), and iNEXT-Discovery (Proposal: 871037). The authors acknowledge the support and the use of resources of Instruct, a Landmark ESFRI project.

## Additional Files

**Additional file 1 — Demonstration of model parameters determination**.

The file contains the mathematical explanations to determine all parameters of the model with deformation.

**Additional file 2 — Excel file containing all parameters of shifts, rotations and deformations applied to phantom data and the parameters found by the algorithm**.

First page is the original parameters, second page corresponds to computed parameters and third page display graphs to compare the parameters.

## Supplementary material

### 1 Update of the model parameters

#### In-plane, image shift: d_*i*_

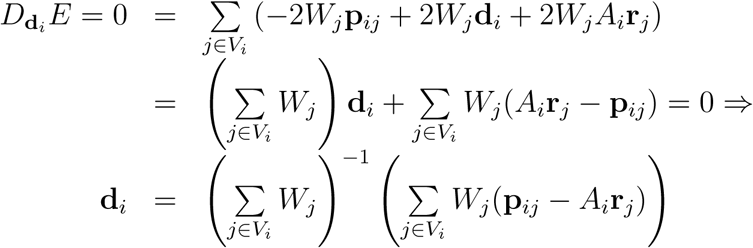

That is, using the current estimates of the alignment parameters we update the in-plane shift as

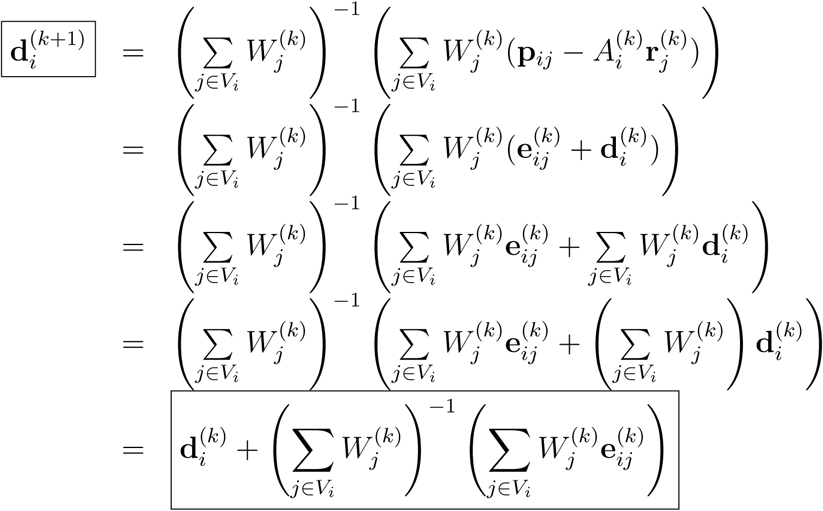

In the particular case of *λ* = 0, then 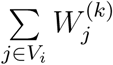 simplifies to *N*_*i*_*I* and the update equation takes the form

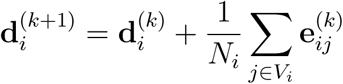

#### 3D landmark: r_*j*_

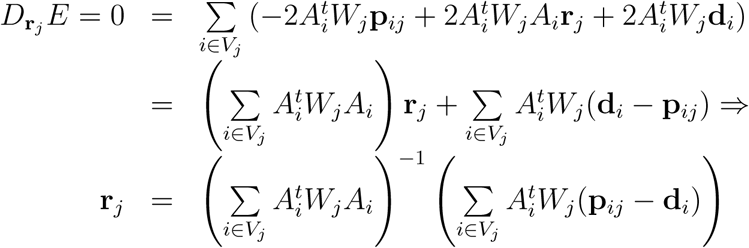

That is, using the current estimates of the alignment parameters we update the 3D landmarks as

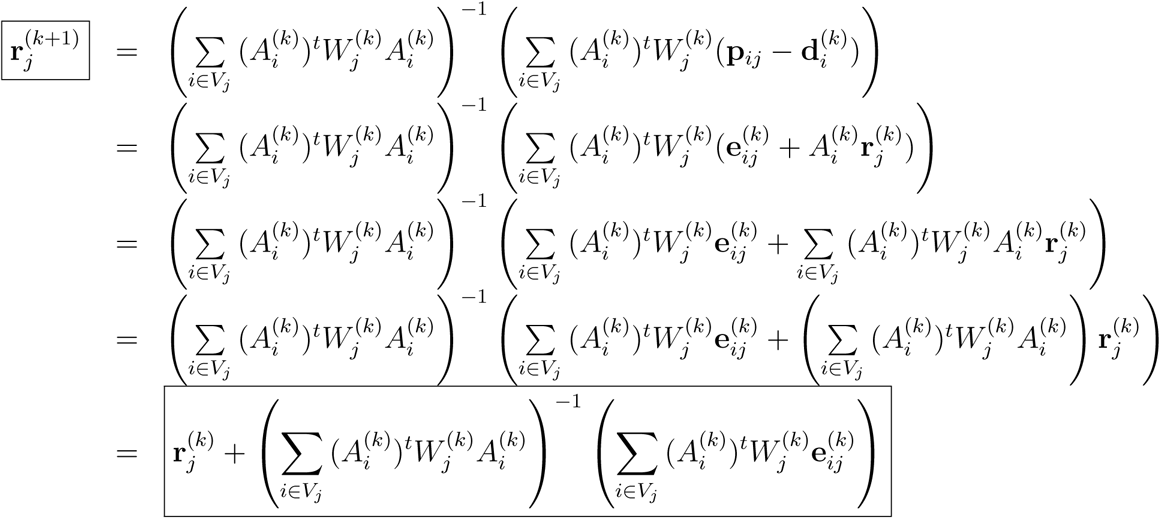

In the case of 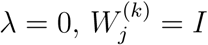.

#### Vertical compression: *t*_*i*_

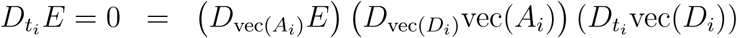

Let us calculate the different terms appearing in the expression above

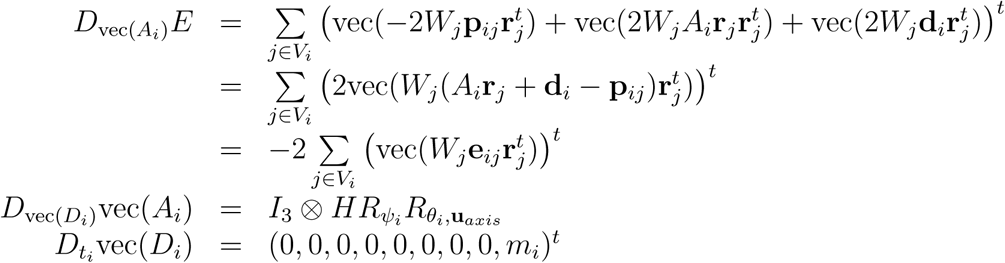

Gathering all terms

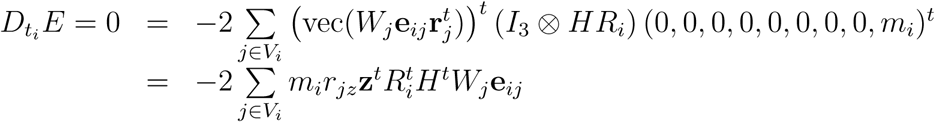

Let us define the scalar product

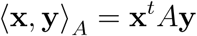

being *A* a symmetric matrix. Then, the derivative above is

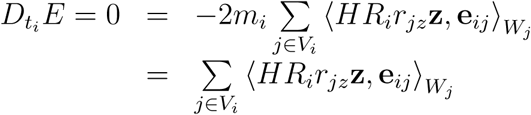

Let us expand **e**_*ij*_ = **p**_*ij*_ − *A*_*i*_**r**_*j*_ − **d**_*i*_ in the expression above

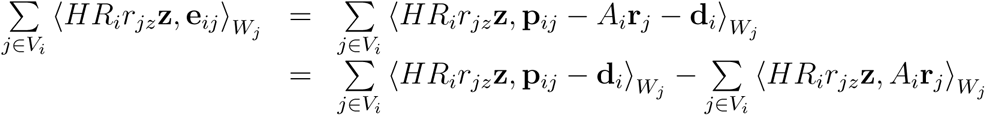

Let us analyze further the term 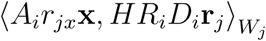

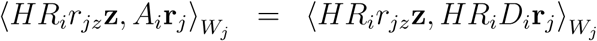

If we decompose *D*_*i*_ as

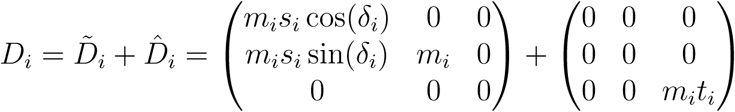

then,

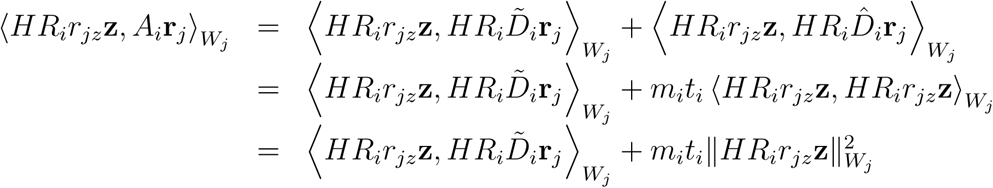

Going back to the derivative

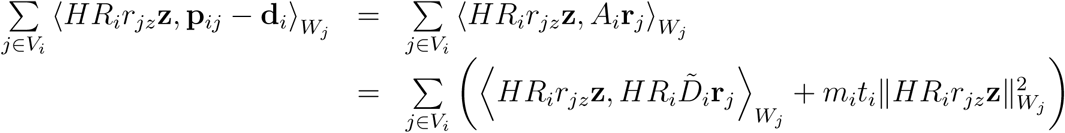

or equivalently

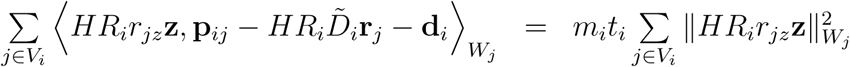

Finally, using the current estimates of the parameters

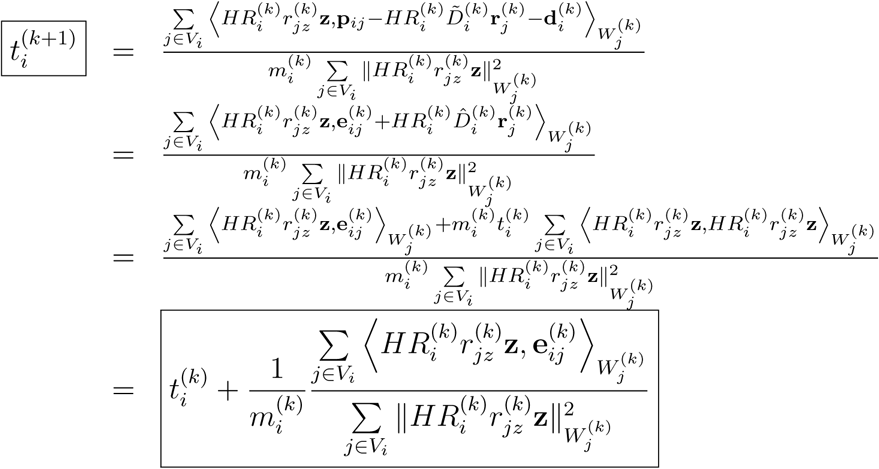

#### Global compression: *m*_*i*_

The derivation for *m*_*i*_ is totally analogous to the one of *t*_*i*_

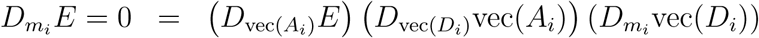

except that

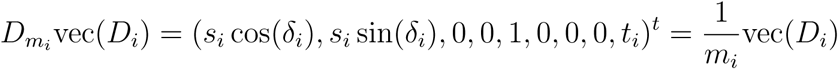

Then

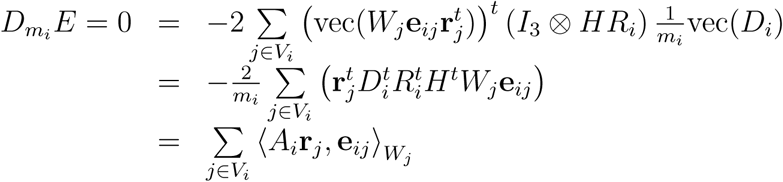

We now expand **e**_*ij*_

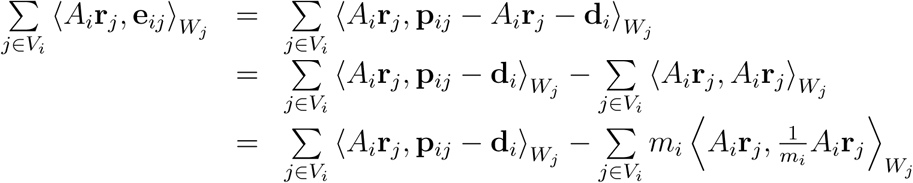

Going back to the derivative equation

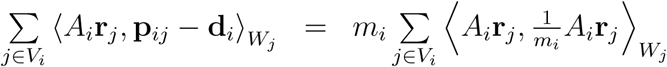

Using the current estimates of the parameters

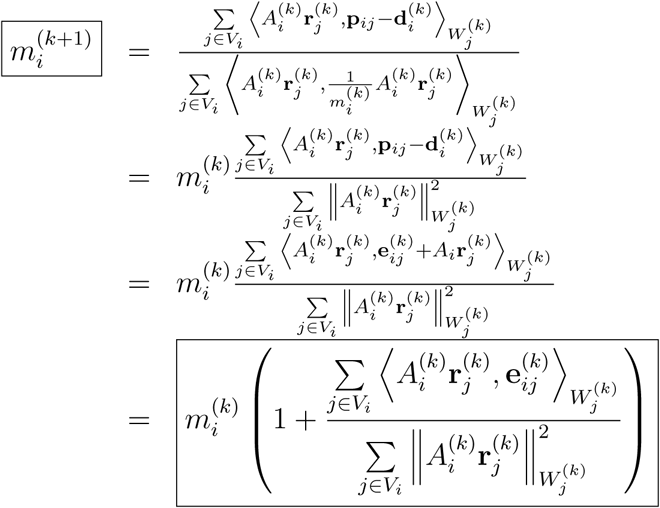

#### Scaling along *X*: *s*_*i*_

Similarly to the previous two cases

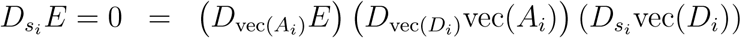

with

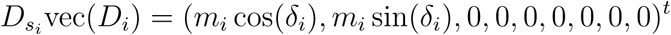

The derivative becomes

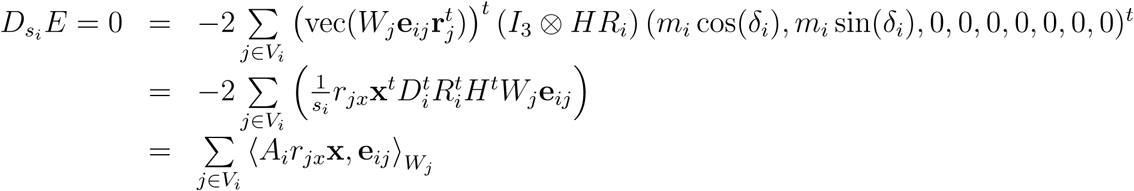

Let us expand **e**_*ij*_

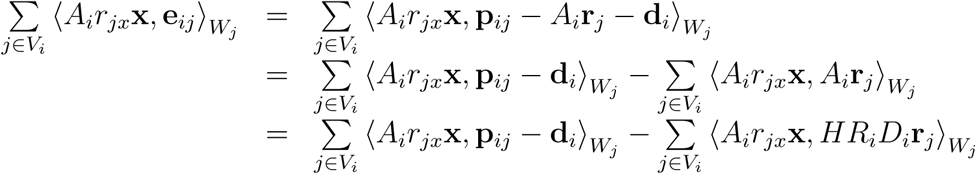

We now decompose *D*_*i*_ as

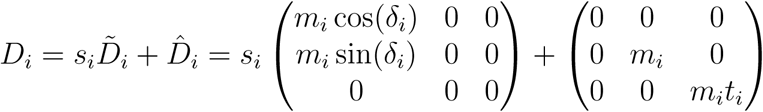

and analyze the term 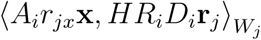

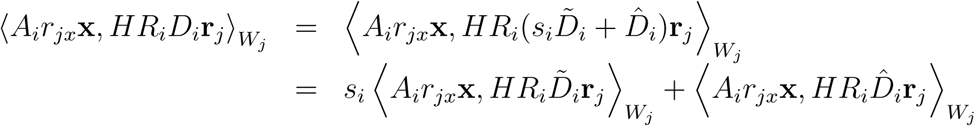

Going back to the derivative

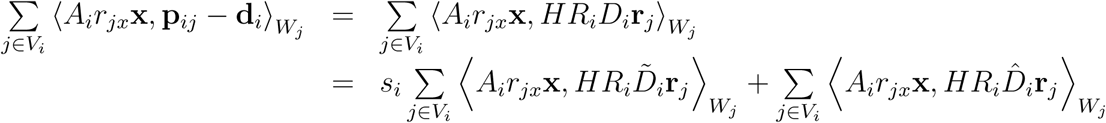

from where

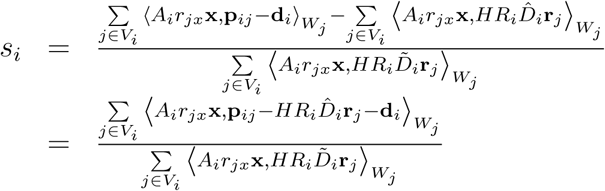

Using the current estimates of the parameters

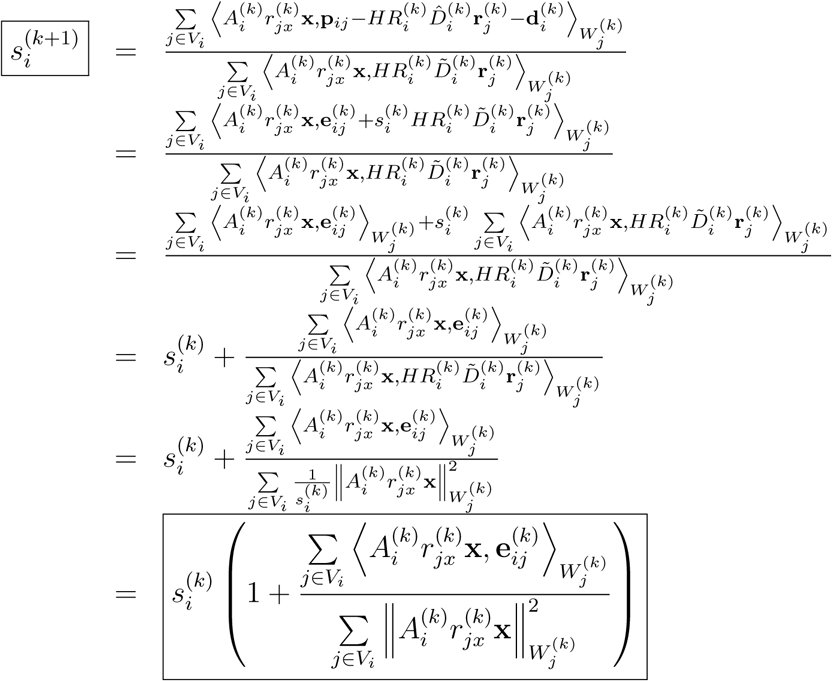

Shearing: *δ*_*i*_ Similarly to the previous cases

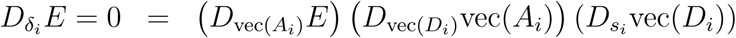

with

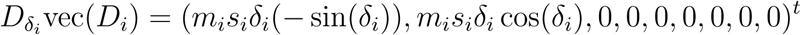

The derivative becomes

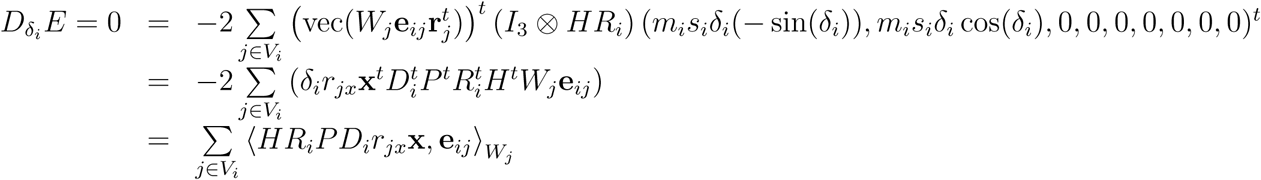

Where 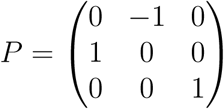. Let’s expand **e**_*ij*_

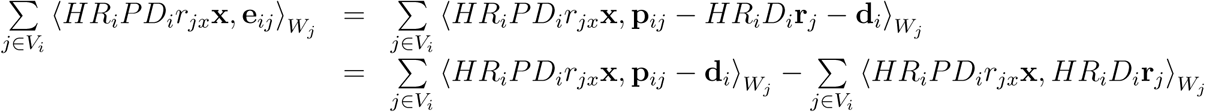

We now decompose *D*_*i*_ as

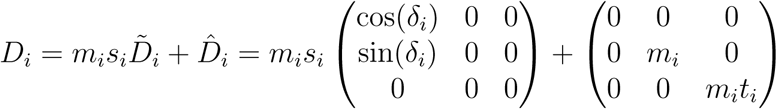

and analyze the term 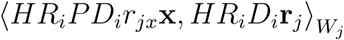

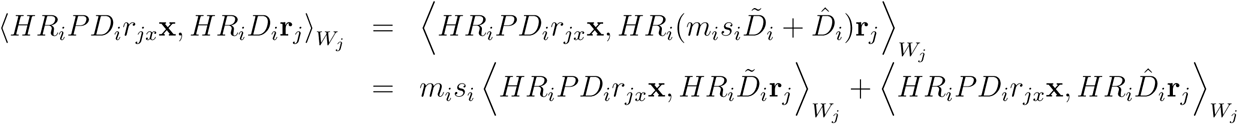

Then

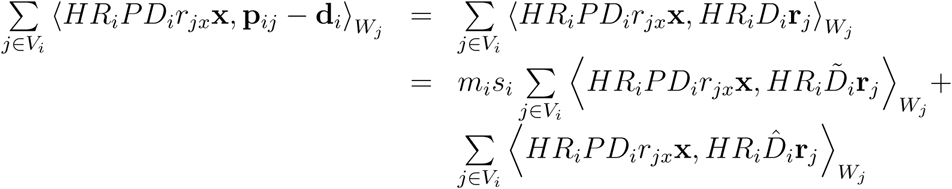

or equivalently

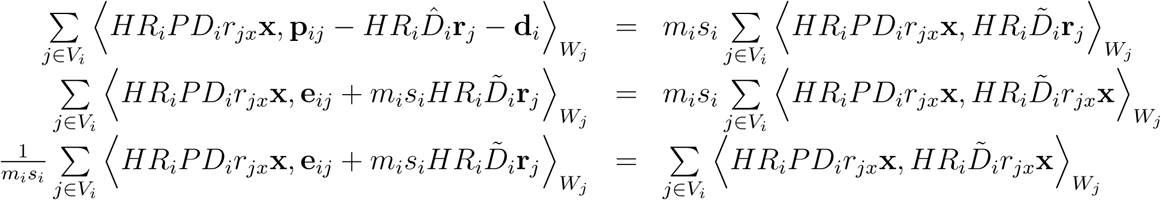

We now decompose as

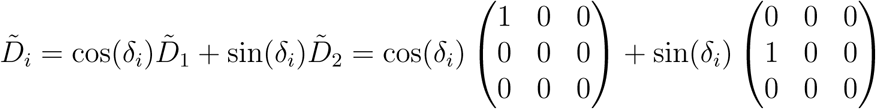

So the right-hand side of the previous expression becomes

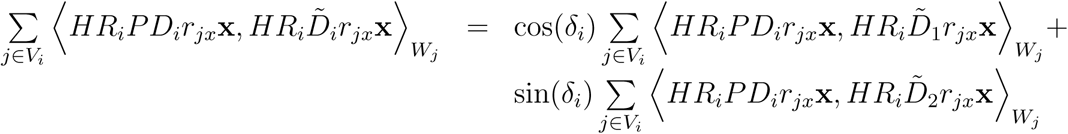

while the left hand side becomes

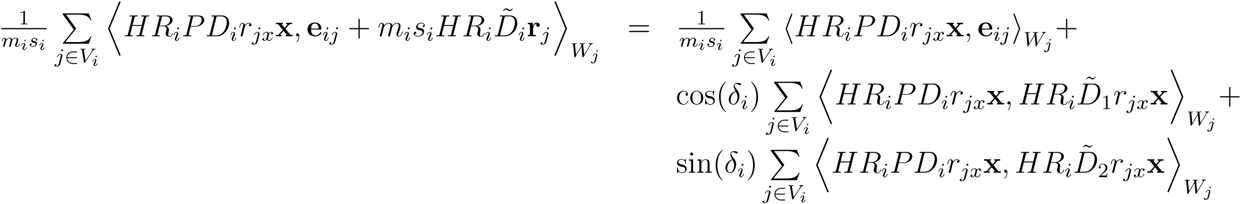

If we call

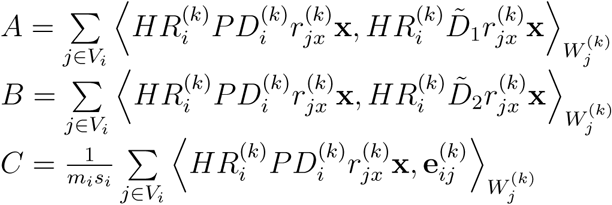

Then, we can update *δ*_*i*_ through the recursion

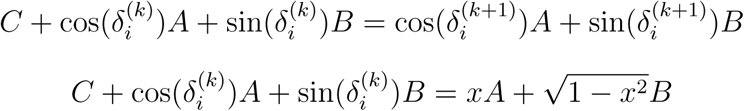

Calling 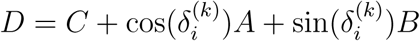

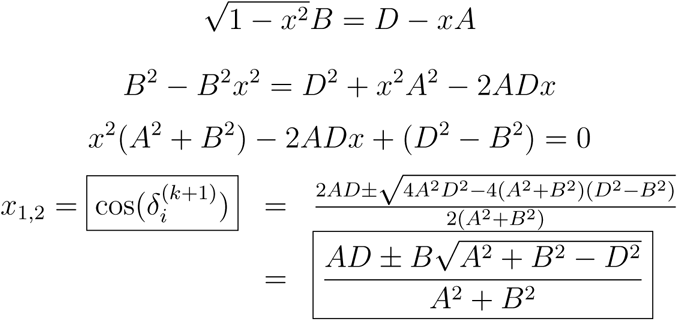

Repeating the same exercise with the sin we would reach

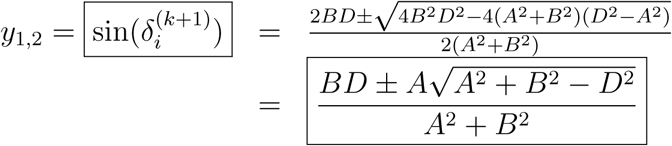

So 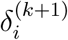 is one of the four angles given by

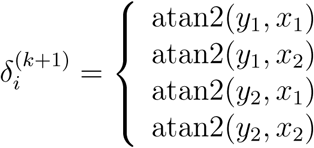

Amongst the fours solutions (which are equivalent from the point of view of the derivative of *E*), we should choose the one that gives smaller *E*.

#### In plane rotation: *ψ*_*i*_

The in-plane rotation matrix is defined as

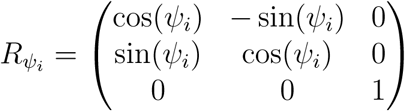

We apply the same methodology as before

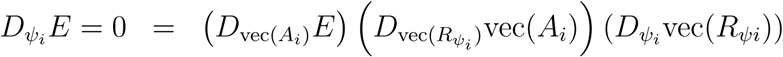

We need to calculate the terms

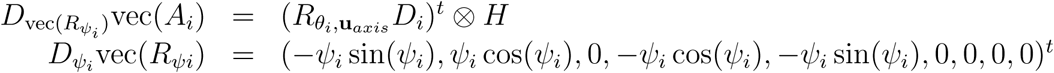

Gathering all terms

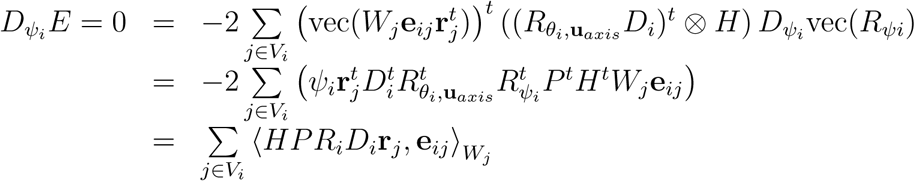

where 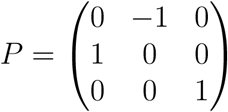.

Let us expand **e**_*ij*_

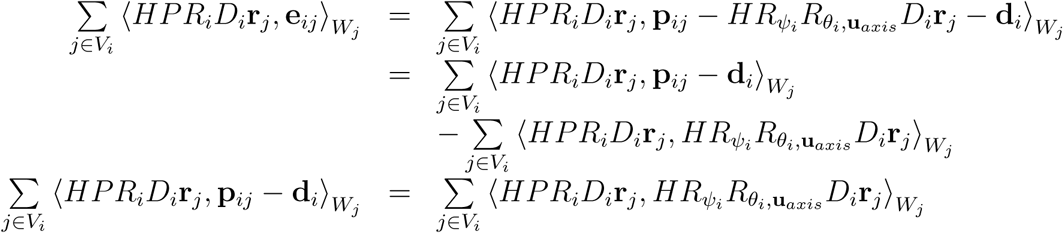

Let us decompose 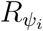 as

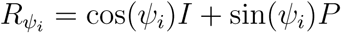

then, the left-hand side of the previous expression becomes

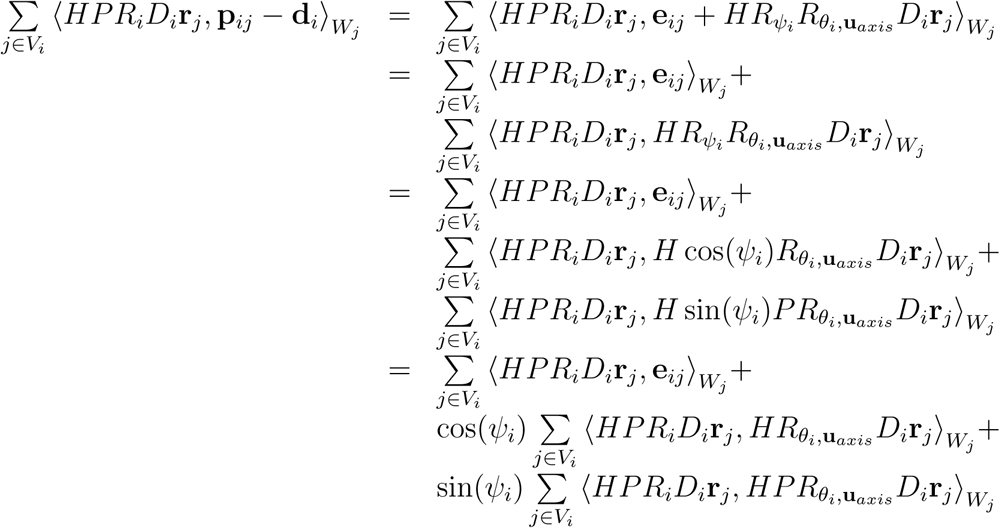

and the right-hand side

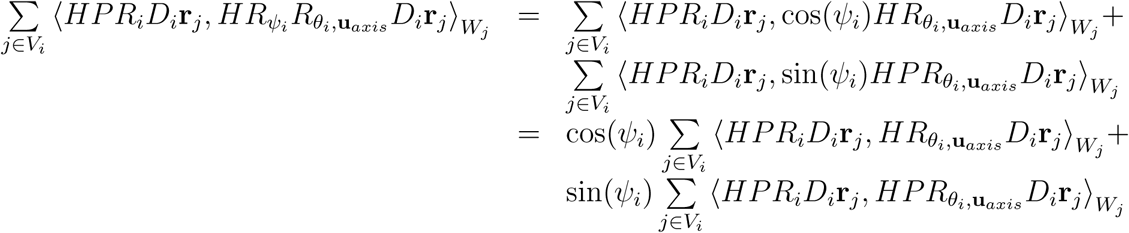

If we call

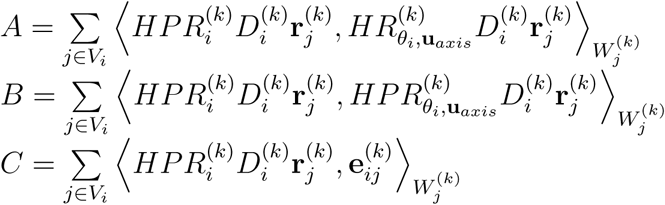

then the in-plane rotation can be updated through the recurrence equation

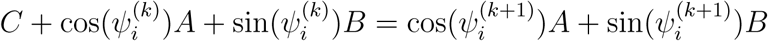

that has the same equation structure as the shearing angle and whose solution is

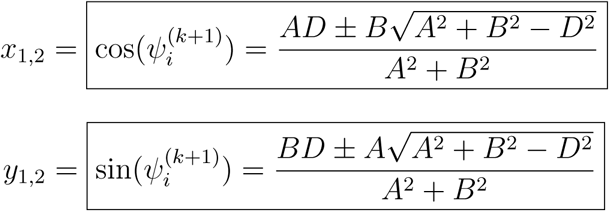

with 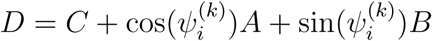.

So 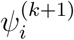 is one of the four angles given by

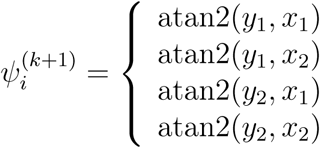

Amongst the fours solutions (which are equivalent from the point of view of the derivative of E), we should choose the one that gives smaller *E*.

#### Tilt axis orientation: *α* and *β*

The arbitrary rotation by an angle *θ*_*i*_ around the axis defined by

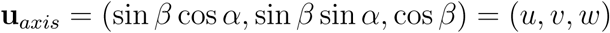

is

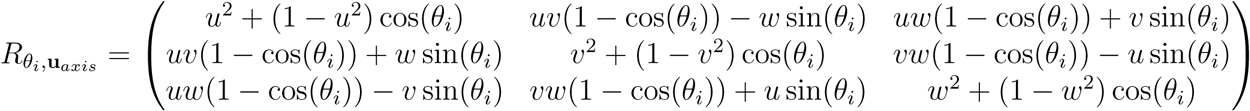

where we have made use of the fact that **u**_*axis*_ is a unitary vector. Although it is feasible to follow the methodology developed along the paper for estimating *α* and *β* from the derivative of *E* with respect to them, the highly nonlinear nature of the equations involved make this path impractical. For this reason, we prefer to employ a global search and a posterior local optimization for estimating these two parameters.

## Appendix

In order to make the article self-contained, let us introduce here the derivative rules we have needed along the article Dattorro (2005)[Appendix D]. Let us define the vectorization operator applied to a matrix *A* as

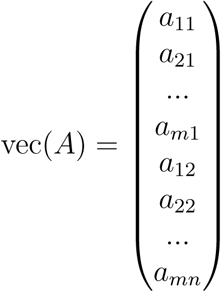

The matrix derivative of a matrix function *F* (whose result is a matrix of size *p* × *q*) with respect to another matrix *X* (of size *m* × *n*) is defined as

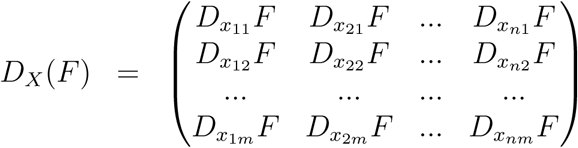

Note that each of the 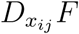 terms is actually another matrix defined as

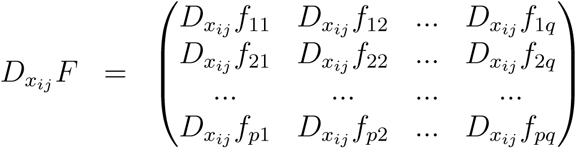

This definition particularizes for a scalar function of a vector as

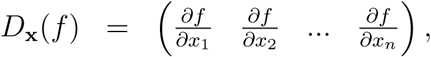

a scalar function of a matrix as

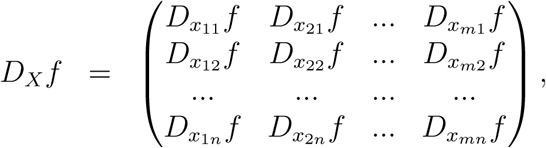

a vector function of a vector as

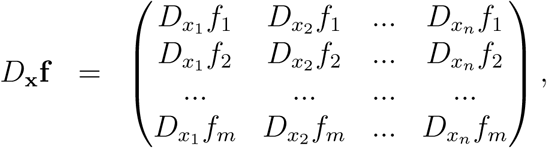

and a vector function of a scalar as

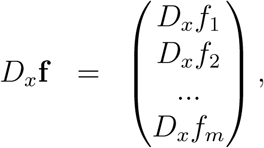

Note that the chain rule does not normally apply with the derivative of a scalar function of a matrix. For this reason, we prefer to arrange this derivative as a vector derivative (for which the chain rule applies), that is,

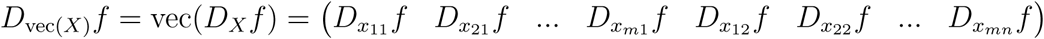

In this way, the chain rule can be written as

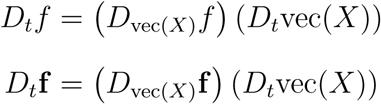

We make use of the following vector and matrix derivatives

- R1: D_*X*_ (a^*t*^Ax) = A^*t*^a
- R2: D_*X*_ (x^*t*^Ab) = Ab
- R3: D_*X*_ (x^*t*^Ax) = (A + A^*t*^)x
- R4: *D*_*X*_(**a**^*t*^*X***b**) = **ab**^*t*^
- R5: *D*_*X*_(**a**^*t*^*X*^*t*^*X***b**) = *X*(**ab**^*t*^ + **ba**^*t*^)
- R6: *D*_*X*_(**a**^*t*^*X*^*t*^*CX***b**) = *C*^*t*^*X***ab**^*t*^ + *CX***ba**^*t*^
- R7: *D*_*X*_(*AXB*) = *B*^*t*^ *⊗ A* where *⊗* denotes the outer product between matrices defined as

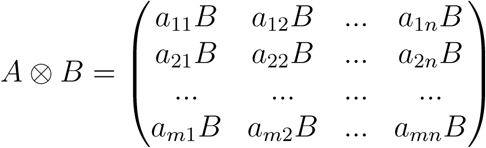
- R8: *D*_*X*_(*AX*) =*I*^*t*^ *⊗ A*
- R9: 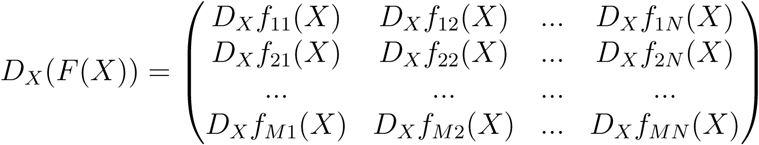 being *F* a matrix function that takes a *K* × *L* matrix *X*, to produce a new matrix of size *M* × *N*. In this case, *D*_*X*_(*F* (*X*)) is a *M* × *N* × *K* × *L* tensor.
- R10: *D*_*X*_(*f* (*g*^*t*^(*X*))) = *D*_*X*_(*g*(*X*))^*t*^*D*_*g*_(*f* (*g*))

